# Alternative stable states, nonlinear behavior, and predictability of microbiome dynamics

**DOI:** 10.1101/2022.08.23.505041

**Authors:** Hiroaki Fujita, Masayuki Ushio, Kenta Suzuki, Masato S. Abe, Masato Yamamichi, Koji Iwayama, Alberto Canarini, Ibuki Hayashi, Keitaro Fukushima, Shinji Fukuda, E. Toby Kiers, Hirokazu Toju

**Author notes:** Correspondence and requests for materials should be addressed to H.F. or H.T.

## Abstract

Microbiome dynamics are both crucial indicators and drivers of human health, agricultural output, and industrial bio-applications. However, predicting microbiome dynamics is notoriously difficult because communities often show abrupt structural changes, such as “dysbiosis” in human microbiomes. We here integrate theoretical and empirical bases for anticipating drastic shifts of microbial communities. We monitored 48 experimental microbiomes for 110 days and observed that various community-level events, including collapse and gradual compositional changes, occurred according to a defined set of environmental conditions. We then confirmed that the abrupt community changes observed through the time-series could be described as shifts between “alternative stable states” or dynamics around complex attractors. Furthermore, collapses of microbiome structure were successfully anticipated by means of the diagnostic threshold defined with the energy landscape analysis of statistical physics or that of a stability index of nonlinear mechanics. These results indicate that abrupt microbiome events in complex microbial communities can be forecasted by extending classic ecological concepts to the scale of species-rich microbial systems.

---

Optimizing biological functions of species-rich systems is a major challenge in both basic and applied sciences^1–7^. Managing the compositions of human gut microbiomes, for example, is essential for preventing diabetes^8,9^, infectious disease^10^, and neuropsychiatric disorders^11^. Likewise, soil and plant-associated microbiomes drive nutrient cycling and pest/pathogen outbreaks in agroecosystems^5,6^, while highly controlled microbiomes facilitate stable and resource-efficient management in biofuel production^7^ and water purification^12^. Nonetheless, it remains generally difficult to control microbial ecosystem functions because species-rich microbial communities often show drastic structural (compositional) changes^13,14^. Thus, predicting such community-scale events remains an essential task.

Drastic changes in biological community structure have been theoretically framed as transient dynamics towards a global equilibrium^15,16^, shifts between alternative equilibria^16,17^, or dynamics around complex forms of attractors^18–20^. Within a state space with a sole equilibrium point, drastic community compositional changes may be observed in the course of convergence to the global equilibrium^15^. In contrast, if multiple equilibria exist within a state space, abrupt community changes can be described as shifts between alternative stable states^17^. In other words, fluctuations in population densities of constituent species (variables) or changes in environments (parameters) can cause shifts of community states from a stable state to the other ones^16,17^. Meanwhile, drastic community changes may be depicted as well in terms of dynamics around periodic/quasi-periodic attractors (i.e., limit cycle or torus) or dynamics around attractors with non-integer dimensions^18,21–23^ (i.e., chaos).

In analyzing empirical time-series data of microbiome structure, these concepts of community dynamics are implemented with two lines of frameworks (Fig. 1a). One is the framework of energy landscape analyses in statistical physics^24–26^, in which stability/instability of possible community states (compositions) are evaluated in the metric of “energy”. In energy landscape analyses, stable states within a state space are defined as community compositions whose energy values are lower than those of adjacent community compositions^24^. Thus, based on the reconstruction of energy landscapes, large community compositional changes are interpreted as transient dynamics towards an equilibrium or shifts between alternative equilibria (Fig. 1a). The other framework for describing abrupt community changes is based on nonlinear mechanics, which allows us to assume the presence of complex forms of attractors^19,20,22,27^. The framework of empirical reconstruction of attractors (“empirical dynamics modeling^28,29^”), in particular, provides a platform for interpreting community dynamics as deterministic processes around any forms of attractors (Fig. 1a). The two frameworks are potentially useful for framing microbial community processes. Nonetheless, it remains to be examined whether drastic changes in microbiome dynamics, such as dysbiosis in human-associated microbiomes^14,30,31^, could be predicted with either or both of the frameworks.

**Fig. 1.**
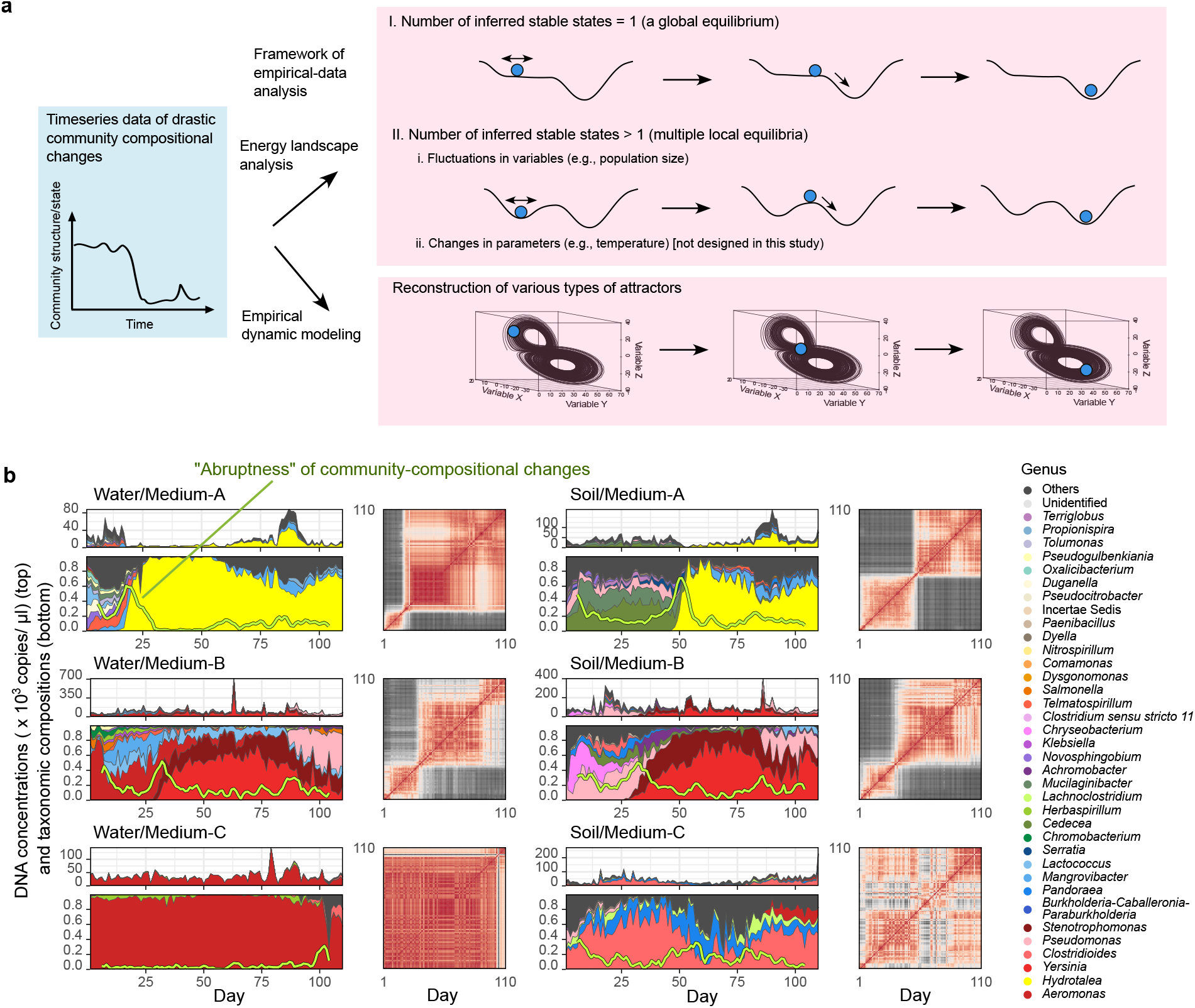
Experimental microbiome dynamics. **a**, Assumptions. Drastic structural changes in microbiome time-series data are interpreted as transient dynamics towards a global equilibrium, shifts between local equilibria (alternative stable states), or dynamics around complex forms of attractors. The former two concepts/models can be examined with an energy landscape analysis and the latter can be explored based on empirical dynamic modeling. **b**, Time-series data of microbial abundance (top left), community compositions (relative abundance; bottom left), and Bray-Curtis dissimilarity (*β*-diversity) of community structure between time points (right) are shown for a representative replicate community of each experimental treatment. The green lines within the relative abundance plots represent the speed and magnitude of community compositional changes (hereafter, “abruptness”) around each target time point (time window = 5 days; time lag = 1 day; see Methods). Note that an abruptness score larger than 0.5 represents turnover of more than 50 % of microbial ASV compositions. See Extended Data Figs. 2-3 for the time-series data of all the 48 communities (8 replicates × 6 treatments).

The major constraint preventing the comparison of the two frameworks is the lack of empirical datasets that simultaneously meet the basic requirements of energy landscape analyses and empirical dynamic modeling. Therefore, by developing a monitoring system of experimental microbiomes, we compile a series of microbiome time-series data with substantial community-compositional changes. By implementing an energy landscape analysis and empirical dynamic modeling, we examine whether the substantial community changes could be anticipated as transient dynamics towards global equilibria, shifts between stable states, or dynamics around complex attractors. Based on the results, we discuss how we can integrate empirical and theoretical studies for predicting and controlling species-rich microbial systems.

## Results

### Experimental microbiome dynamics

To obtain time-series datasets of diverse microbiome dynamics, we constructed six types of microbiomes based on the combinations of two inoculum sources (soil and pond water microbiomes; hereafter, Soil and Water) and three media differing in chemical properties (oatmeal, oatmeal-peptone, and peptone; hereafter, Medium-A, B, and C, respectively), each with eight replicates (Extended Data Fig. 1). We kept the experimental system at a constant temperature condition and sampled a fraction of each microbiome and added fresh media every 24 hours for 110 days. For each of the six experimental treatment, 880 community samples were obtained (in total, 110 time points × 8 replicates × 6 treatments = 5,280 community samples), providing rich information for exploring stable states of community structure by means of energy landscape analyses. In total, the dataset represented population dynamics of 264 prokaryote amplicon sequence variants (ASVs) belonging to 108 genera. Using quantitative amplicon sequencing^32^ for estimating 16S ribosomal RNA gene (16S rRNA) copy concentrations of respective microbes in each microbiome, we determined the dynamics of both “relative” and “absolute” ASV abundance (Fig. 1b; Extended Data Figs. 1-3). By estimating not only relative but also absolute abundance, we were able to reconstruct respective ASVs’ population dynamics (increase/decrease), satisfying the requirements for applying empirical dynamic modeling^19,20,22^.

The experimental microbiomes exhibited various types of dynamics depending on source inocula and culture media (Fig. 1b; Extended Data Figs. 2-3). Specifically, sharp decline of taxonomic diversity^33^ and abrupt (sudden and substantial) community structural changes (see “abruptness” index in Fig. 1b) were observed in Water/Medium-A, Soil/Medium-A, and Water/Medium-B treatments (abruptness > 0.5). Within these treatments, taxonomic compositions and timing of abrupt shifts in community structure varied among replicate communities (Extended Data Fig. 3). Large shifts of community compositions through time were observed as well in Soil/Medium-B treatment, albeit the community shifts were more gradual (maximum abruptness through time-series, 0.36 ∼ 0.57; Extended Data Fig. 3). In contrast, Medium-C condition yielded relatively steady microbiome dynamics with continuously low taxonomic diversity (e.g., dominance of *Aeromonas* in Water/Medium-C treatment), although shifts of dominant taxa were observed latter in the experiment in some replicate communities (Extended Data Fig. 3). In all the six treatments, the *α*-diversity (Shannon diversity) of ASVs displayed fluctuations, but not monotonic decrease, through time (Extended Data Fig. 1e).

### Framework 1: energy landscape analysis

By compiling the microbiome time-series data, we examined the distributions of stable states within the multidimensional space of community structure based on an energy landscape analysis^24^. Because no variation in environmental conditions was introduced through the time-series in our experiment, a fixed “energy landscape” of community states was assumed for each of the six treatments. On this assumption, shifts between alternative stable states are attributed to perturbations to variables (i.e., population density of microbial ASVs) but not to “regime shifts^34–36^”, which, by definition, requires energy landscape reorganization caused by changes in environmental parameters (i.e., temperature).

In each experimental treatment, multiple stable states were estimated to exist (Fig. 2; Extended Data Fig. 4), indicating that the observed abrupt changes in community compositions could be described as shifts between alternative stable states. Therefore, in this approach of statistical physics^24–26^, community dynamics are divided into phases of fluctuations around local equilibrium points and those of shifts into adjacent equilibria. In other words, the presence of multiple equilibrium points (Extended Data Fig. 4), by definition, means that the observed dynamics of the experimental microbiomes are not described as transient dynamics towards a sole equilibrium point.

**Fig. 2.**
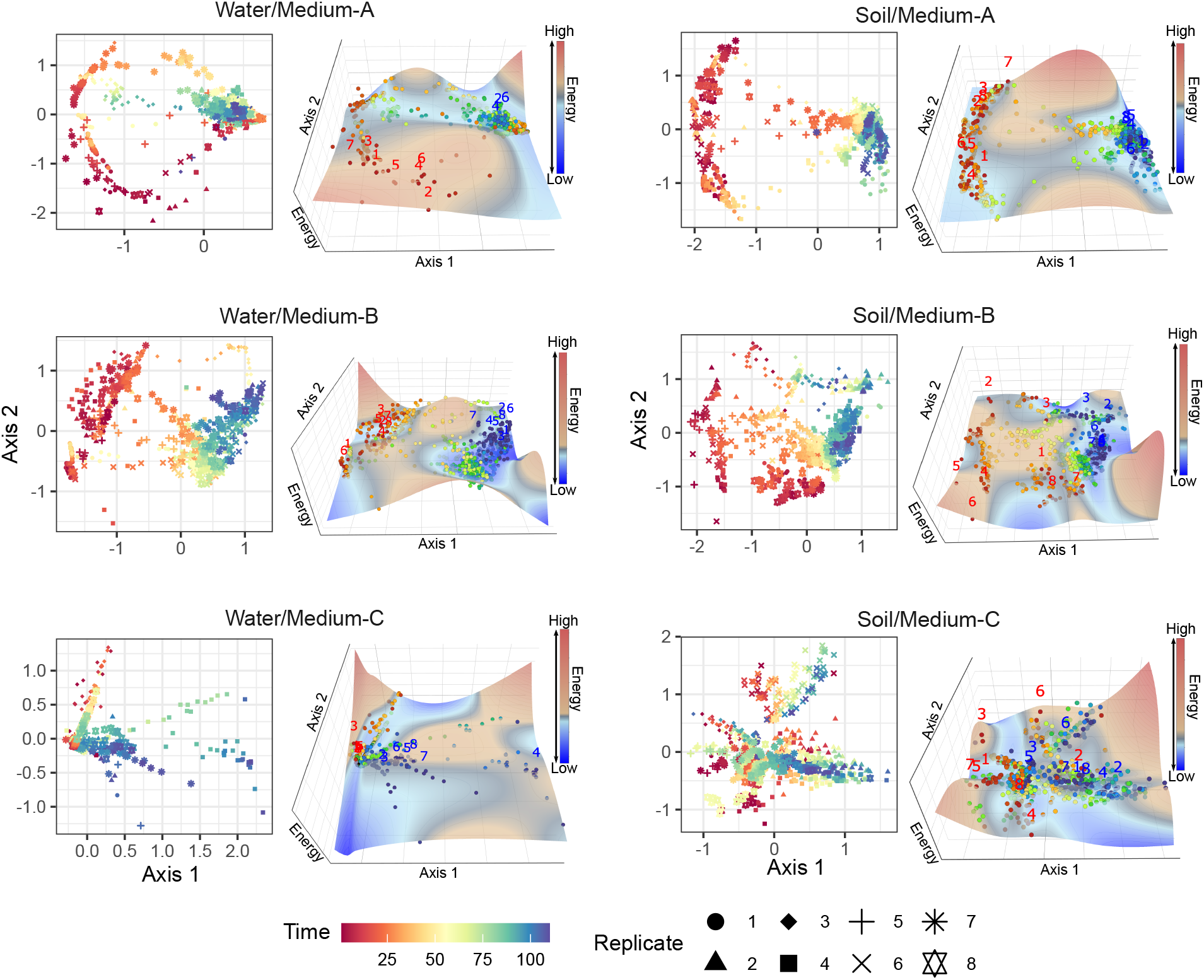
Energy landscapes of community structure. The community structure of respective time points on NMDS axes (left) and reconstructed energy landscape on the NMDS surface (right) are shown for each experimental treatment. Community states (ASV compositions) located at lower-energy regions are inferred to be more stable on the energy landscapes. The shapes of the landscapes were inferred based on a smoothing spline method with optimized penalty parameters. On the energy landscapes, community states of Day 1 and Day 110 are respectively shown in red and blue numbers representing replicate communities.

### Framework 2: empirical dynamic modeling

We next analyzed the time-series data based on the framework of empirical dynamic modeling. We first focused on the population dynamics (increase/decrease) of the microbial ASVs constituting the microbial communities. In ecology, population dynamics data have often been analyzed with methods assuming linear dynamics (i.e., without considering “state dependency^37^”). Meanwhile, a series of empirical dynamics modeling approaches applicable to nonlinear time-series processes, such as simplex projection^20^ and sequential locally weighted global linear maps^19^ (S-map), have been increasingly adopted to capture key properties lost with linear dynamic assumptions (Fig. 3a). We found that ca. 85 % of the microbial populations in our experiments exhibited nonlinear behavior (i.e., nonlinearity parameter *θ* > 0; Fig. 3b). This result suggests the predominance of nonlinear dynamics over linear dynamics in microbial populations^32^, in line with populations of other organismal groups such as fish^38^ and plankton^21^.

**Fig. 3.**
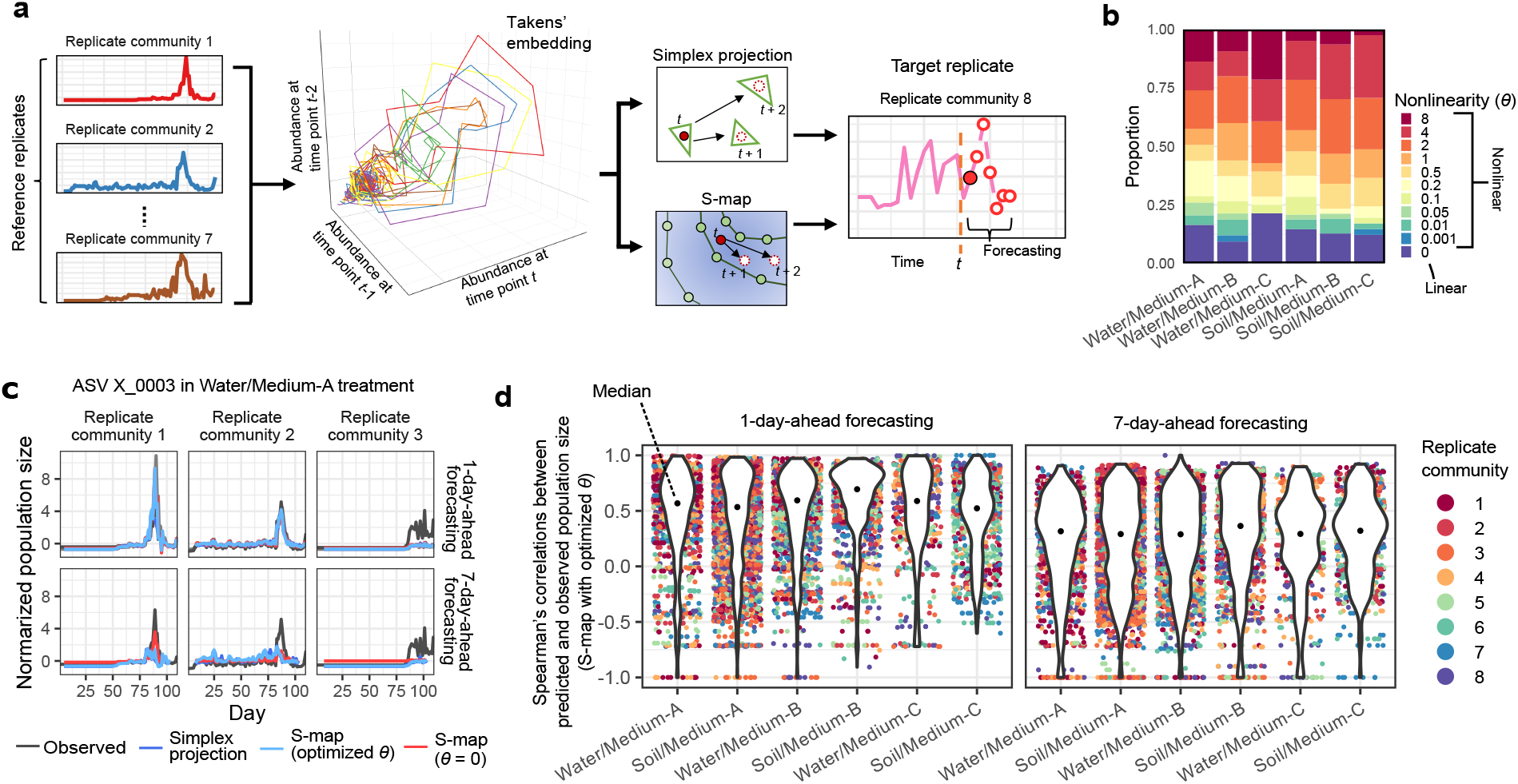
Forecasting population-level dynamics based on attractor reconstruction. **a**, Workflow of forecasting. For a target replicate community, the reference database of state space is reconstructed based on the time-series data of other replicate communities (i.e., reference replicate communities). Future abundance of each ASV in a target replicate community was then predicted using the state-space reference databases (see Methods for details). **b**, Nonlinearity parameters (*θ*). Proportions of microbial ASVs showing linear (*θ* = 0) and nonlinear (*θ* > 0) population dynamics are shown. **c**, Example of population-level forecasting. Predicted and observed abundance through the time-series are shown for simplex projection, S-map with optimized nonlinearity parameter (optimized *θ*), and S-map assuming linearity (*θ* = 0). For each target replicate community, the remaining seven replicate communities were used as references. Results are shown for one-day-ahead and seven-day-ahead forecasting of an ASV (X_0003) in replicate nos. 1-3 of Water/Medium-A treatment. See Extended Data Fig. 5 for detailed results. d, Spearman’s correlation between predicted and observed population size is shown for each microbial ASV in each replicate community.

We then reconstructed the attractors of nonlinear dynamics based on Takens’ embedding theorem^39^ (Fig. 3a). To examine the performance of the attractor reconstruction, we conducted forecasting of the population dynamics of respective microbial ASVs by means of simplex projection and S-map (Fig. 3c). The population density (16S rRNA copy concentration) of an ASV in a target replicate community at time point *t* + *p* (*p* represents time steps in forecasting) was forecasted based on the ASV’s population density at time point *t* and time-series data of other replicate communities (hereafter, reference replicate communities; see Methods for details; Fig. 3a). For many microbial ASVs, predicted population densities was positively correlated with observed ones (Fig. 3c-d; Extended Data Fig. 5). As expected, correlation between predicted and observed population size increased with increasing number of reference replicate communities, suggesting dependence of forecasting skill on the size of state-space reference databases (Extended Data Fig. 6).

By assembling the forecasting results of respective ASVs at the community level, we further conducted forecasting of microbiome compositions (Fig. 4a; Extended Data Fig. 7). The forecasting precision of community-level dynamics varied depending on culture media and the dissimilarity (*β*-diversity) of community structure between target and reference replicates (Fig. 4b). Despite the utility of the forecasting platform, we observed high prediction error immediately after the peak of abrupt community changes (Fig. 4c; Extended Data Fig. 8). Although the nonlinear method (S-map with optimized *θ*) captured the observed abrupt shifts of community compositions within a narrower time window than the linear method (S-map with *θ* = 0) (Fig. 4c), quantitative forecasting of abrupt community changes seemed inherently difficult.

**Fig. 4.**
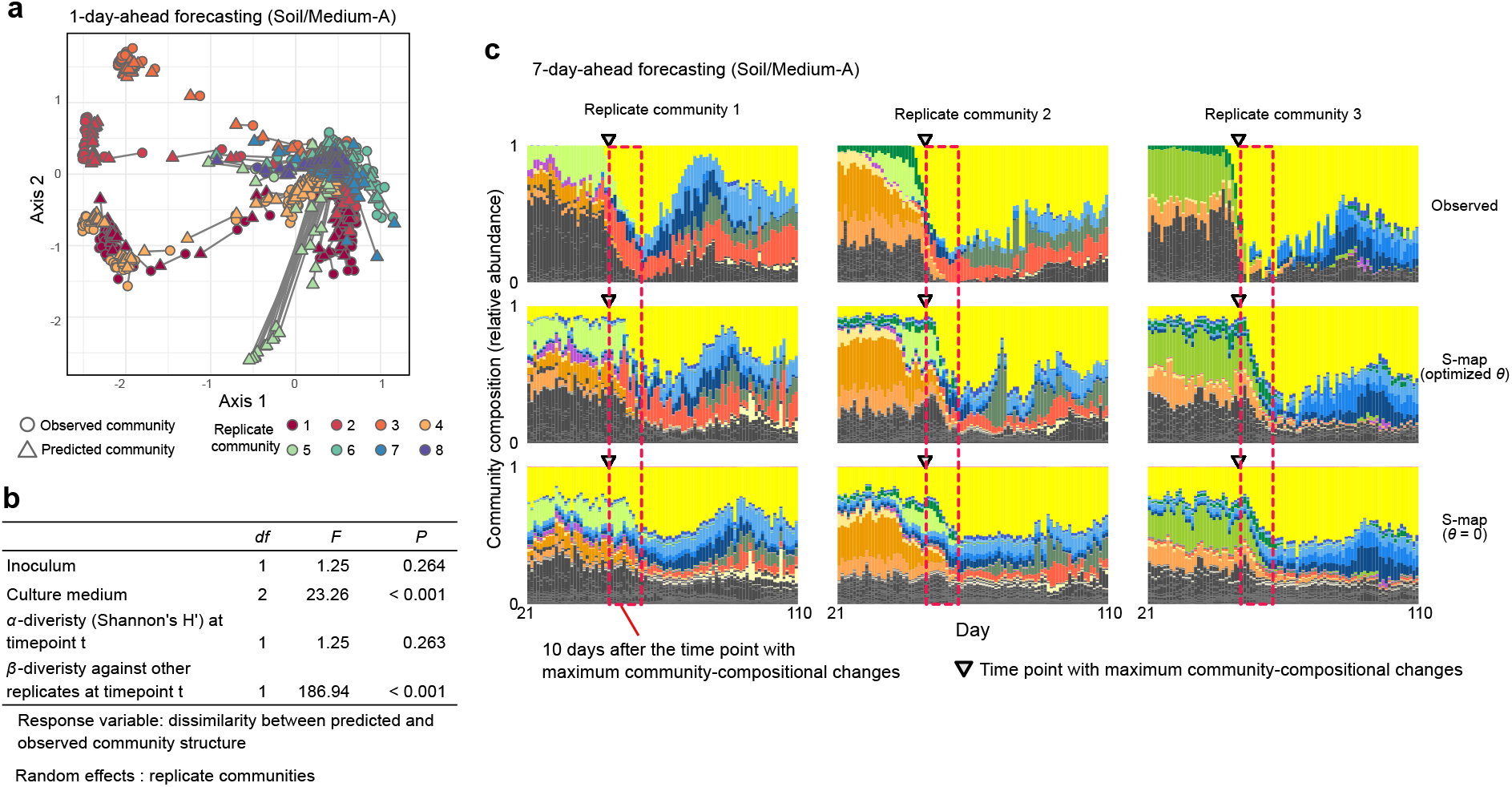
Forecasting community-level dynamics based on attractor reconstruction. **a**, Community-level forecasting. Predicted and observed community structure is linked for each day on the axes of NMDS (prediction based on S-map with optimized *θ*; one-day-ahead forecasting). Results of Soil/Medium-A treatment are shown: see Extended Data Fig. 7 for full results). **b**, Factors explaining variation in community-level prediction results. A generalized linear mixed morel (GLMM) of dissimilarity between predicted and observed community structure was constructed (one-day-ahead forecasting). **c**, Detailed comparison of nonlinear and linear forecasting approaches. S-map results with optimized nonlinearity parameter were compared with results of S-map assuming linear dynamics for all ASVs.

Nonetheless, even if precise forecasting of community compositional dynamics remains challenging, prediction of the occurrence of abrupt community changes *per se* may be possible. Thus, we next examined whether potential of abrupt community changes could be evaluated through microbiome dynamics.

### Anticipating abrupt community shifts

Based on the frameworks of the energy landscape analysis and empirical dynamic modeling, we explored ways for anticipating abrupt events in community dynamics. In the former framework roach, the reconstructed energy landscapes were used to estimate “energy gap” and “stable-state entropy” indices, which represented stability/instability of community states^24^ (Fig. 5a). In the latter framework, the inferred Jacobian matrices of the multi-species time-series dynamics (see Methods) were used to calculate “local Lyapunov stability^40^” and “local structural stability^41^”. We examined how these indices could help us forecast large community-compositional shifts such as those observed in Medium-A and Medium-B treatments (Fig. 1b).

**Fig. 5.**
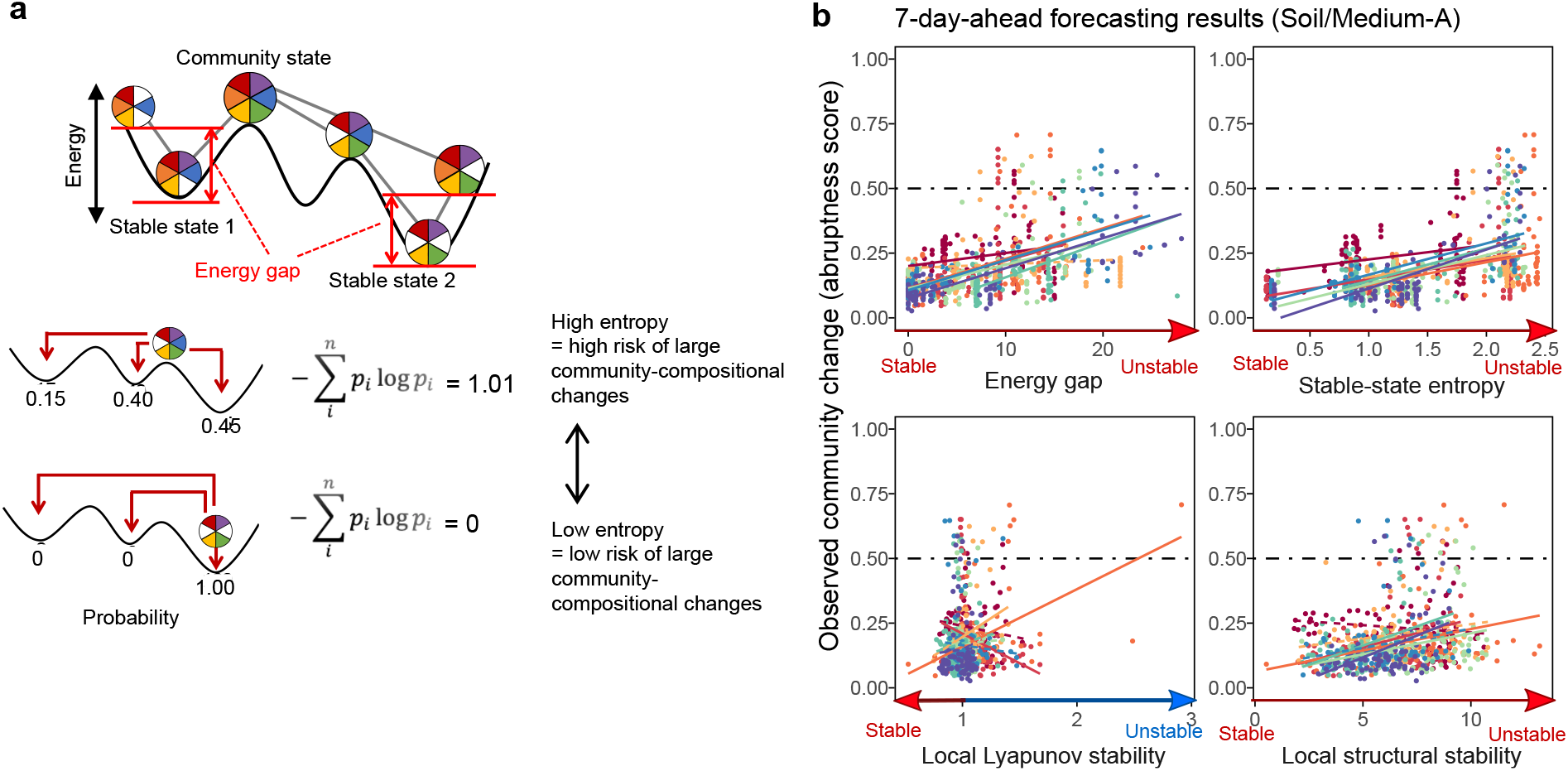
Anticipating abrupt community shifts. **a**, Energy gap index. In the framework of the energy landscape analysis, difference between the energy of a current community state from that of the local energy minimum is defined as the “energy gap” index for anticipating drastic community changes. **b**, Stable-state entropy index. Shannon’s entropy estimates based on random-walk simulations towards alternative stable states are expected to represent instability of current community states on energy landscapes (see Methods for details). **c**, Relationship between the degree of community-compositional changes (abruptness) and each signal index. Note that a high score of energy gap, stable-state entropy, or local Lyapunov/structural stability potentially represents an unstable state. Significant/non-significant regressions within respective replicates are shown with solid/dashed lines for each panel [false discovery rate (FDR)]. See Extended Data Fig. 9 for detailed results.

Among the signal indices examined, energy gap or stable-state entropy of community states (Fig. 5a) was significantly correlated with the degree of future community changes in Medium-A and Medium-B treatments (FDR < 0.05 for all treatments; Fig. 5b; Extended Data Fig. 9). In the seven-day-ahead forecasting of abrupt community-compositional changes (abruptness > 0.5), for example, stable-state entropy showed relatively high diagnostic performance on the two-dimensional surface of detection rate (sensitivity) and false detection rate (1 – specificity) as represented by receiver operating characteristic (ROC) curve^42^. Specifically, although the small number of points with abruptness greater than 0.5 (Extended Data Fig. 10) precluded the application of the ROC analysis in Soil/Medium-B treatment, diagnostic performance as evaluated by area under the ROC curve (AUC) ranged from 0.726 to 0.957 in other Medium-A and Medium-B treatments (Fig. 6a).

**Fig. 6.**
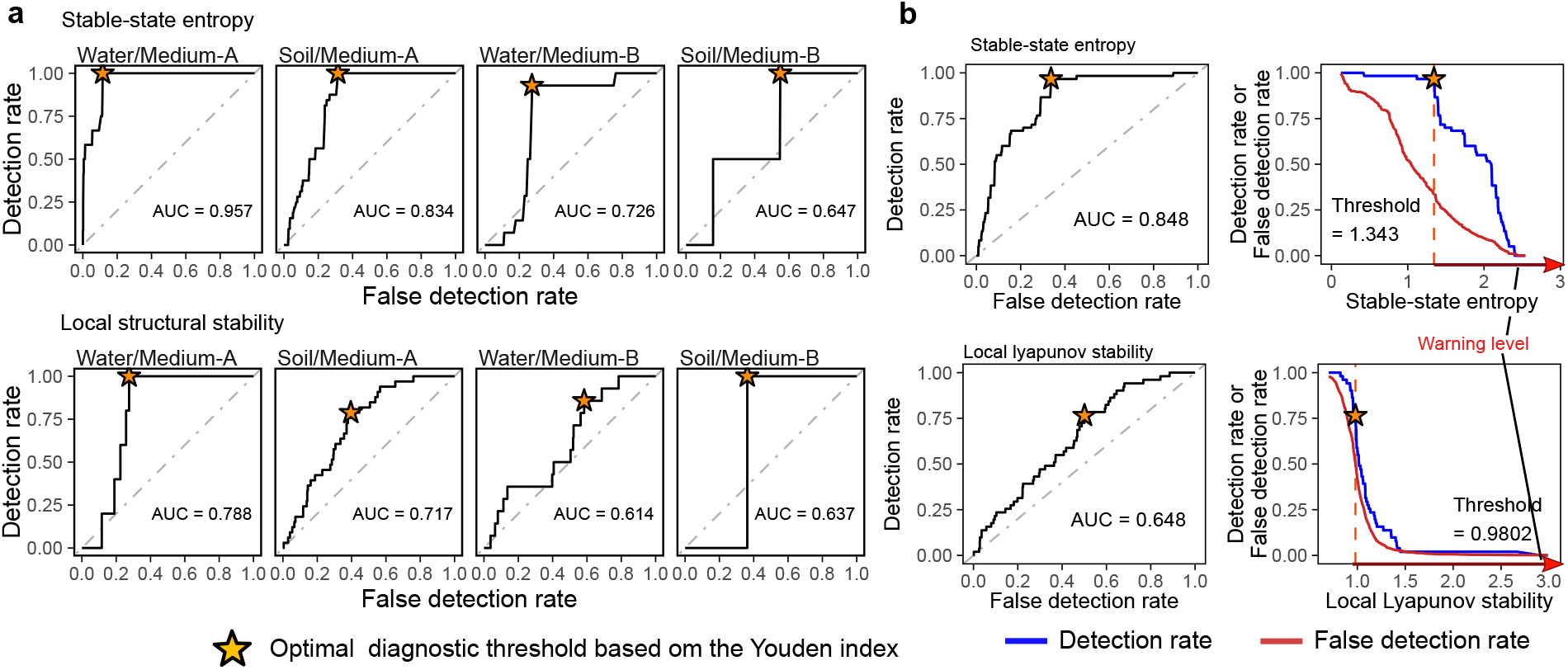
Thresholds for anticipating drastic community events. **a**, ROC analysis of diagnostic performance. On the two-dimensional surface of detection- and false-detection rates of abrupt community-compositional changes (abruptness > 0.5), area under the ROC curve (AUC) and optimal detection rate (asterisk) were calculated for each warning signals (seven-day-ahead forecasting). A high AUC value indicates a high detection rate of abrupt community events with a relatively low false detection rate (maximum AUC value = 1). Note that the low AUC values may be attributed to the small number of points with abruptness > 0.5 in Soil/Medium-B treatment. See Extended Data Fig. 10 for full results. **b**, Optimal thresholds for anticipating community collapse. For stable-state entropy (top) and local Lyapunov stability (bottom), diagnostic threshold for warning abrupt community changes was obtained based on the Youden index after compiling all the data of Medium-A and Medium-B treatments.

Local Lyapunov or structural stability was correlated with the degree of community changes as well, but the correlations were less consistent among experimental treatments than energy gap and stable-state entropy (Fig. 5b; Extended Data Fig. 9). Meanwhile, local structural stability exhibited exceptionally high diagnostic performance in Water/Medium-A treatment (AUC = 0.788; Fig. 6a; Extended Data Fig. 10). Thus, local Lyapunov or structural stability can be used as signs of future microbiome collapse, although further technical improvement in the state space reconstruction of species-rich communities (e.g., multi-view distance regularized S-map^43^) may be needed to gain consistent forecasting performance across various types of microbiomes.

By further utilizing the frameworks of the energy landscape analysis and empirical dynamic modeling, we next examined the availability of diagnostic thresholds for anticipating community collapse. For this aim, we first focused on stable-state entropy because its absolute values in the unit of well-known entropy index (Fig. 5a) were comparable across diverse types of biological communities. Based on the ROC curve representing all the stable-state entropy data of Medium-A and Medium-B treatments, the balance between detection and false-detection rates were optimized with the Youden index^42^. With a relatively high AUC score (0.848), the threshold stable-state entropy was set as 1.343 (Fig. 6b). In the same way, we calculated the threshold value for local Lyapunov stability because this index originally had a tipping value (= 1) for diagnosing community-level stability/instability^40^. Indeed, the estimated threshold of local Lyapunov stability on the ROC curve was 0.9802, close to the theoretically expected value (Fig. 6b).

## Discussion

By compiling datasets of experimental microbiome dynamics under various environmental (medium) conditions, we here tested whether two lines of ecological concepts could allow us to anticipate drastic compositional changes in microbial communities. Despite decades-long discussion on alternative stable/transient states of community structure^15–17,35,36^, the application of the concept to empirical data of species-rich communities has been made feasible only recently with the computationally intensive approach of statistical physics (energy landscape analyses^24^). On the other hand, the concept of dynamics around complex forms of attractors has been applicable with the emerging framework of nonlinear mechanics^27,40,41^ (e.g., empirical dynamic modeling), microbiome time-series data satisfying the requirements of the analytical frameworks remained scarce^32^. Thus, this study, which was designed to apply both frameworks, provided a novel opportunity for fuel feedback between empirical studies of species-rich communities and theoretical studies based on classic/emerging ecological concepts.

Our analysis showed drastic events in microbiome dynamics, such as those observed in dysbiosis of human-gut microbiomes^13,14^, could be forecasted, at least to some extent, by framing microbiome time-series data as shifts between alternative stable states or dynamics around complex attractors. In the forecasting of abrupt community changes observed in our experimental microbiomes, the former concept (model) seemingly outperformed the latter (Figs. 5-6). This result is of particular interest, because concepts or models more efficiently capturing dynamics of empirical data are expected to provide more plausible planforms in not only prediction but also control of biological community processes. Nonetheless, given the ongoing methodological improvements of nonlinear mechanics frameworks for describing empirical time-series data^43^, further empirical studies comparing the two concepts are necessary.

A key next step for forecasting and controlling microbial (and non-microbial) community dynamics is to examine whether common diagnostic thresholds could be used to anticipate collapse of community structure. This study provided the first empirical example that the tipping value theoretically defined in noncolinear mechanics^40^ (local Lyapunov stability = 1) could be actually used as a threshold for alerting microbiome collapse. Likewise, although estimates of diagnostic thresholds can vary depending on the definition of community collapse (e.g., abruptness > 0.5 in this study), stable-state entropy scores greater than 1.3 can be used to anticipate undesirable community events (dysbiosis) across medical, agricultural, and industrial applications.

Given that changes in environmental parameters were not incorporated into our experimental design, it remains another important challenge to reveal how distributions of stable states or forms of attractors are continually reshaped by changes in environmental parameters through community dynamics^17,34,35^. Experimental manipulation of “external” environmental parameters in microcosms, for example, will expand the target of research into microbiome systems potentially driven by regime shifts^34–36^. Likewise, environmental alternations caused by organisms *per se*^44–46^ deserve further investigations as potential drivers of drastic community shifts.

Controlling biological functions at the ecosystem level is one of the major scientific challenges in the 21^st^ century^5,47,48^. Interdisciplinary approaches that further integrate microbiology, ecology, and mathematics are becoming indispensable for maximizing and stabilizing microbiome-level functions, and for providing novel solutions to a broad range of humanity issues spanning from human health to sustainable industry and food production.

## Methods

### Continuous-culture of microbiome

To set up experimental bacterial communities, we prepared two types of source inocula (soil and aquatic microbiomes) and three media (oatmeal, oatmeal-peptone, and peptone): for each combination of source media and inocula (six experimental treatments), eight replicate communities were established (in total, two source microbiomes × three media × eight replicates = 48 experimental communities; Extended Data Fig. 1a). We used natural microbial communities including diverse species, rather than “synthetic” communities with pre-defined diversity, as source microbiomes of the experiment. One of the source microbiomes derives from the soil collected from the A layer (0-10 cm in depth) in the research forest of Center for Ecological Research, Kyoto University, Kyoto, Japan (34.972 °N; 135.958 °E). After sampling, 600 g of the soil was sieved with a 2-mm mesh and then 5 g of the sieved soil was mixed in 30 mL autoclaved distilled water. The source microbiome was further diluted 10 times with autoclaved distilled water. The source aquatic microbiome was prepared by collecting 200 mL of water from a pond (“Shoubuike”) near Center for Ecological Research (34.974 °N, 135.966 °E). In the laboratory, 3 mL of the collected water was mixed with 27 mL of distilled water in a 50 mL centrifuge tube. We then introduced the source soil or aquatic microbiomes into three types of media: oatmeal broth [0.5% (w/v) milled oatmeal (Nisshoku Oats; Nippon Food Manufacturer); Medium-A], oatmeal-peptone broth [0.5% (w/v) milled oatmeal + 0.5% (w/v) peptone (Bacto Peptone; BD; lot number: 7100982); Medium-B], and peptone broth [0.5% (w/v) peptone; Medium-C]. In our preliminary experiments, microbiomes cultured with Medium-A (oatmeal) tended to show high species diversity, while those cultured with Medium-C (peptone) were constituted by smaller number of bacterial species. Thus, we expected that diverse types of microbiome dynamics would be observed with the three medium conditions. Among the three media, Medium-B had the highest concentrations of non-purgeable organic carbon (NPOC) and total nitrogen (TN), while Medium-A was the poorest both in NPOC and TN: Medium-C had the intermediate properties (Extended Data Fig. 1b).

In each well a of 2-mL deep well plate, 200 µL of a diluted source microbiome and 800 µL of medium were installed. The deep-well plate was kept shaken at 1,000 rpm using a microplate mixer NS-4P (AS ONE Corporation, Osaka) at 23 °C for five days. After the five-day pre-incubation, 200 µL out of 1,000-µL culture medium was sampled from each of the 48 deep wells after mixing (pipetting) every 24 hours for 110 days. In each sampling event, 200 µL of fresh medium was added to each well so that the total culture volume was kept constant. In total, 5,280 samples (48 communities/day × 110 days) were collected. Note that on Day 82, 200-µL of fresh Medium-B was accidentally added to samples of Medium-A but not to those of Medium-B. While the microbiomes under Medium-A treatments experienced increase in total DNA copy concentrations late in the time-series, relative abundance remained relatively constant from Day 60 to 110 (Extended Data Figs. 2-3), suggesting limited impacts of the accidental addition of the medium on microbial community compositions.

To extract DNA from each sample, 25 µL of the collected aliquot was mixed with 50 µL lysis buffer (0.0025 % SDS, 20 mM Tris (pH 8.0), 2.5 mM EDTA, and 0.4 M NaCl) and proteinase K (×1/100). The mixed solution was incubated at 37 °C for 60 min followed by 95 °C for 10 min and then the solution was vortexed for 10 min to increase DNA yield.

### Quantitative 16S rRNA sequencing

To reveal the increase/decrease of population size for each microbial ASV, a quantitative amplicon sequencing method^32,49^ was used based on Illumina sequencing platform. While most existing microbiome studies were designed to reveal the “relative” abundance of microbial ASVs or operational taxonomic units (OTUs), analyses of population dynamics, in principle, require the time-series information of “absolute” abundance. In our quantitative amplicon sequencing, five standard DNA sequence variants with different concentrations of artificial 16S rRNA sequences (0.1, 0.05, 0.02, 0.01, and 0.005 nM) were added to PCR master mix solutions (Extended Data Fig. 1a). The DNA copy concentration gradient of the standard DNA variants yielded calibration curves between Illumina sequencing read numbers and DNA copy numbers (concentrations) of the 16S rRNA region in PCR reactions, allowing estimation of original DNA concentrations of target samples^32,49^ (Extended Data Fig. 1c-d). The five standard DNAs were designed to be amplified with a primer set of 515f^50^ and 806rB^51^ but not to be aligned to the V4 region of any existing prokaryote 16S rRNA. Note that the number of 16S rRNA copies per genome generally varies among prokaryotic taxa^52^ and hence 16S rRNA copy concentration is not directly the optimal proxy of cell or biomass concentration. However, in our study, estimates of 16S rRNA copy concentrations are used to monitor increase/decrease of abundance (i.e., population dynamics) *within* the time-series of each microbial ASV: hence, variation in the number 16S rRNA copy numbers among microbial taxa had no qualitative effects on the subsequent population- and community-ecological analyses. Even if the concentrations of PCR inhibitor molecules in DNA extracts vary among time-series samples, potential bias caused by such inhibitors can be corrected based on the abovementioned method using internal standards (i.e., standard DNAs within PCR master solutions).

Prokaryote 16S rRNA region was PCR-amplified with the forward primer 515f fused with 3–6-mer Ns for improved Illumina sequencing quality and the forward Illumina sequencing primer (5’-TCG TCG GCA GCG TCA GAT GTG TAT AAG AGA CAG- [3–6-mer Ns] – [515f] -3’) and the reverse primer 806rB fused with 3–6-mer Ns for improved Illumina sequencing quality^53^ and the reverse sequencing primer (5’-GTC TCG TGG GCT CGG AGA TGT GTA TAA GAG ACA G [3–6-mer Ns] - [806rB] -3’) (0.2 µM each). The buffer and polymerase system of KOD One (Toyobo) was used with the temperature profile of 35 cycles at 98 °C for 10 s, 55 °C for 30 s, 68 °C for 50 s. To prevent generation of chimeric sequences, the ramp rate through the thermal cycles was set to 1 °C/sec^54^. Illumina sequencing adaptors were then added to respective samples in the supplemental PCR using the forward fusion primers consisting of the P5 Illumina adaptor, 8-mer indexes for sample identification^55^ and a partial sequence of the sequencing primer (5’-AAT GAT ACG GCG ACC ACC GAG ATC TAC AC - [8-mer index] - TCG TCG GCA GCG TC -3’) and the reverse fusion primers consisting of the P7 adaptor, 8-mer indexes, and a partial sequence of the sequencing primer (5’-CAA GCA GAA GAC GGC ATA CGA GAT - [8-mer index] - GTC TCG TGG GCT CGG -3’). KOD One was used with a temperature profile: followed by 8 cycles at 98 °C for 10 s, 55 °C for 30 s, 68 °C for 50 s (ramp rate = 1 °C/s). The PCR amplicons of the samples were then pooled after a purification/equalization process with the AMPureXP Kit (Beckman Coulter). Primer dimers, which were shorter than 200 bp, were removed from the pooled library by supplemental purification with AMpureXP: the ratio of AMPureXP reagent to the pooled library was set to 0.6 (v/v) in this process. The sequencing libraries were processed in an Illumina MiSeq sequencer (271 forward (R1) and 231 reverse (R4) cycles; 10% PhiX spike-in).

### Bioinformatics

In total, 67,537,480 sequencing reads were obtained in the Illumina sequencing. The raw sequencing data were converted into FASTQ files using the program bcl2fastq 1.8.4 distributed by Illumina. The output FASTQ files were then demultiplexed with the program Claident v0.2. 2018.05.29^56^. The sequencing reads were subsequently processed with the program DADA2^57^ v.1.13.0 of R 3.6.0 to remove low-quality data. The molecular identification of the obtained ASVs was performed based on the naive Bayesian classifier method^58^ with the SILVA v.132 database^59^. In total, 399 prokaryote (bacterial or archaeal) ASVs were detected. We obtained a sample × ASV matrix, in which a cell entry depicted the concentration of 16S rRNA copies of an ASV in a sample. In this process of estimating original DNA copy numbers (concentrations) of respective ASVs from sequencing read numbers in each sample, the samples in which Pearson’s coefficients of correlations between sequencing read numbers and standard DNA copy numbers (i.e., correlation coefficients representing calibration curves) were less than 0.7 (in total, 430 samples out of 5,280 samples) were removed as those with unreliable estimates. Samples with less than 350 reads were discarded as well. Because missing values within time-series data are not tolerated in some of the downstream analyses (e.g., empirical dynamic modeling), they were substituted by interpolated values, which were obtained as means of the time points immediately before and after focal missing time points. The ASVs that appeared 5 or more samples in any of the replicate communities were retained in the following analyses: 264 ASVs representing 108 genera remained in the dataset. From the sample × ASV matrix, we calculated *α*-diversity (Shannon’s *H′*) of the ASV compositions in each experimental replicate on each day. We also evaluated dissimilarity of community compositions in all pairs of sampling days in each replicate community using Bray-Curtis metric of *β*-diversity as implemented in the vegan 2.5.5 package^60^ of R. For each ASV in each replicate community, a parameter representing the nonlinearity of the population dynamics^19,38^ (*θ*) was estimated based on S-map analysis of absolute abundance as detailed below in order to evaluate the overall nature of the time-series data.

### Community dynamics

We evaluated the degree of community-compositional changes for time point *t* based on the Bray-Curtis *β*-diversity through time. To remove effects of minor fluctuations and track only fundamental changes of community structure, average community compositions from time points *t* – 4 to *t* and those from *t* + *p* to *t* + *p* + 4 (i.e., 5-day time-windows) were calculated before evaluating degree of community changes for time point *t* and time step *p* in each replicate community. Dissimilarity of community compositions between the time windows before (from *t* – 4 to *t*) and after (*t* + *p* to *t* + *p* + 4) each target time point *t* with a given time lag *p* was calculated based on Bray-Curtis *β*-diversity as a measure of abrupt (sudden and substantial) community changes (hereafter, “abruptness” of community-compositional changes). A high value of this index indicates that abrupt community-compositional changes occurred around time point *t*, while a low value represents a (quasi-)stable mode of community dynamics. We also evaluated temporal changes of community compositions using nonmetric multidimensional scaling (NMDS) with the R package vegan.

### Energy landscape analysis

On the assumption that drastic changes in microbiome dynamics are described as shifts between local equilibria (i.e., alternative stable states), we reconstructed the structure of the “energy landscape^24,25^” in each experimental treatment (tutorials of energy landscape analyses are available at https://community.wolfram.com/groups/-/m/t/2358581). Because external environmental conditions (e.g., temperature) was kept constant in the experiment, a fixed “energy landscape” of community states was assumed for each of the six experimental treatments. Therefore, probabilities of community states 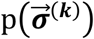 are given by

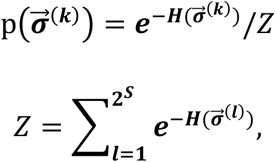

where 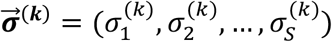 is a community state vector of *k*-th community state and *S* is the total number of taxa (e.g., ASVs, species, or genera) examined. 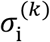 is a binary variable that represents presence (1) or absence (0) of taxon *i*: i.e., there are a total of 2^*S*^community states. Then, the energy of the community state 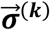 is given by

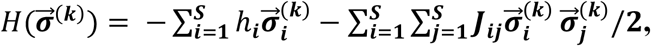

where *h*_*i*_ is the net effect of implicit abiotic factors, by which *i*-th taxon is more likely to present (*h*_*i*_ > 0) or not (*h*_*i*_ < 0), and *J*_*ij*_ represents a co-occurrence pattern of *i*-th and *j*-th taxa. Since the logarithm of the probability of a community state is inversely proportional to 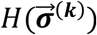, a community state having lower *H* is more frequently observed. To consider dynamics on an assembly graph defined as a network whose 2^*S*^nodes represent possible community states and the edges represents transition path between them (two community states are adjacent only if they have the opposite presence/absence status for just one species), we assigned energy to nodes with the above equation, and so imposed the directionality in state transitions. Then, we identified the stable state communities as the energy minima of the weighted network (nodes having the lowest energy compared to all its neighbors), and determined their basins of attraction based on a steepest descent procedure starting from each node. The data of ASV-level compositions were used in the calculation of community state energy using Mathematica v12.0.0.0. The “energy” estimates were then plotted against the NMDS axes representing community states of the microbiome samples in each experimental treatment. Spline smoothing of the landscape was performed with optimized penalty scores using the mgcv v.1.8-40 package^61^ of R.

### Empirical dynamic modeling

In parallel with the energy landscape analysis assuming the presence of local equilibria, we also analyzed the microbiome time-series data by assuming the presence of complex attractors. In this aim, we applied the framework of “empirical dynamic modeling^19,20,29,40^”. In general, biological community dynamics are driven by a number of variables (e.g., abundance of respective species and abiotic environmental factors). In the analysis of a multi-variable dynamic system in which only some of variables are observable, state space constituted by time-lag axes of observable variables can represent the whole system as shown in Takens’ embedding theorem^39^. Thus, for each ASV in each experimental treatment, we conducted Takens’ embedding to reconstruct state space which consisted of time-delayed coordinates of the ASV’s absolute abundance (e.g., 16S rRNA copy concentration estimates). The optimal number of embedding dimensions^29,39^ (*E*) was obtained by finding *E* giving the smallest root-mean-square error (RMSE) in pre-run forecasting with simplex projection^20^ or S-map^19^ as detailed below. Taking into account a previous study examining embedding dimensions^62^, optimal *E* was explored within the range from 1 to 20. Prior to the embedding, all the variables were *z*-standardized (i.e., zero-mean and unit-variance) across the time-series of each ASV in each replicate community.

### Population-level forecasting

Based on the state space reconstructed with Takens’ embedding, simplex projection^20^ was applied for forecasting of ecological processes in our experimental microbiomes. For each target replicate community, univariate embedding of each ASV was performed using the data of the seven remaining replicate communities. Therefore, the reference databases for the embedding did not include the information of the target replicate community (Fig. 2a), providing platforms for evaluating forecasting skill.

In simplex projection, a coordinate within the reconstructed state space was explored at a focal time point (*t*^*\**^) within the population dynamics of a focal ASV in a target replicate community (e.g., replicate community 8): the coordinate can be described as [*x*_*target_rep*_(*t*^*\**^), *x*_*target_rep*_(*t*^*\**^ – 1), *x*_*target_rep*_(*t*^*\**^ – 2)] when *E* = 3. For the focal coordinate, *E* + 1 neighboring points are explored from the reference database consisting of the seven remaining replicate communities (e.g., replicate communities 1–7; Fig. 2a). For each of the neighboring points, the corresponding points at *p*-time-step forward (*p*-days ahead) are identified. The abundance estimate of a focal ASV in the target replicate community at *p*-time-step forward [e.g., *x*_*target_rep*_(*t*^*\**^ + *p*)] is then obtained as weighted average of the values of the highlighted *p*-time-step-forward points within the reference database (Fig. 2a). The weighting was decreased with Euclidean distance between *x*_*target_rep*_(*t*^*\**^) and its neighboring points within the reference database. This forecasting of population dynamics was performed for each ASV in each target replicate community at each time point. The number of time steps in the forecasting (i.e., *p*) was set at 1 (one-day-ahead forecasting) and 7 (seven-day-ahead forecasting).

While simplex projection explores neighboring points around a target point, S-map^19^ uses all the data points after weighting contributions of each point within a reference database using a parameter representing nonlinearity of the system. In Takens’ embedding, the state space of a target replicate for forecasting at time *t* is defined as

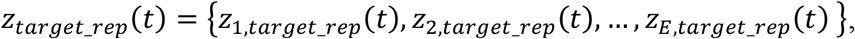

where *E* is embedding dimension. Values on the second and higher dimensions {*z*_2,*target*_*rep*_ (*t*), …, *z*_*E, target*_*rep*_ (*t*)} are represented by time-delayed coordinates of a focal ASV. Likewise, the state space of the remaining replicates (i.e., the reference database) is defined as

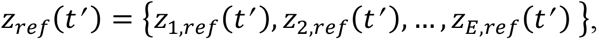

where *t′* represents each of non-overlapping time points within the reference database [e.g., {10001, 10002, …, 10110} and {20001, 20002, …, 20110} for reference replicate 1 and 2, respectively]. For a target time point *t*^*^ within the time-series data of a target replicate community, a local linear model ***C*** is produced to predict the future abundance of a focal ASV [i.e., *z*_1, *target*_*rep*_ (*t*^*^ ± *p*)] from the state-space vector at time point *z*_*target*_*rep*_ (*t*^*^)as follows:

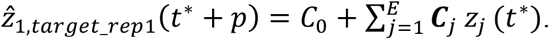

This linear model is fit to the vectors in the reference databases. In the regression analysis, data points close to the target point *z*_*target*_*rep*_ (*t*^*^) have greater weighting. The model ***C*** is then the singular value decomposition solution to the equation *b* = *A****C***. In this equation, *b* is set as an *n*-dimensional vector of the weighted future values of *z*_1,*ref*_ (*t*_*i*_*′*) for each point (*t*_*i*_*′*) in the reference database (*n* is the number of points in the set of *t*_*i*_*′*):

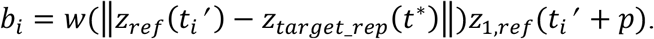

Meanwhile, *A* is an *n* × *E* dimensional matrix given by

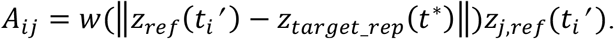

The weighting function *w* is defined as

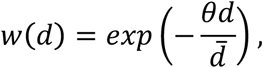

where *θ* is the parameter representing the nonlinearity of the data, while mean Euclidean distance between reference database points and the target point in the target experimental replicate is defined as follows:

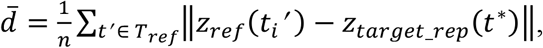

where *T*_*ref*_ denotes the set of *t*_*i*_*′*. In our analysis, the optimal value of *θ* was explored among eleven levels from 0 (linearity) and 8 (strong nonlinearity) for each ASV in each target replicate based on the RMSE of forecasting (optimal *θ* was selected among 0, 0.001, 0.01, 0.05, 0.1, 0.2, 0.5, 1, 2, 4, and 8). The local linear model ***C*** was estimated for each time point in the target replicate data.

We then performed direct comparison between linear and nonlinear approaches of forecasting based on empirical dynamics modeling. Specifically, to assume linear dynamics in S-map method, the nonlinearity parameter *θ* was set 0 for all the ASVs. We then compared forecasting results between linear (*θ =* 0) and nonlinear (optimized *θ*) approaches. For the forecasting of ASVs in a target replicate community, the data of the remaining seven communities (reference databases) were used as mentioned above.

For each ASV in each of the 48 experimental replicates, Spearman’s correlation coefficients between predicted and observed abundance (16S rRNA copy concentrations) were calculated for each of the nonlinear/linear forecasting methods [simplex projection, S-map with optimized *θ*, and S-map assuming linearity (*θ* = 0)]. We also examined null model assuming no change in community structure for a given time step. The time points (samples) excluded in the data-quality filtering process (see Bioinformatics sub-section) were excluded from the above evaluation of forecasting skill.

### Reference database size and forecasting skill

To evaluate potential dependence of forecasting skill on the size of reference databases, we performed a series of analyses with varying numbers of reference replicate communities. For replicate community for a target replicate community, a fixed number (from 1 to 7) of other replicate communities within each experimental treatment were retrieved as reference databases: all combinations of reference communities were examined for each target replicate community. For each microbial ASV in each target replicate community, forecasting of population size was performed based on S-map with optimized *θ* as detailed above. Spearman’s correlation between predicted and observed population size across the time-series was then calculated for each ASV in each target replicate community. The correlation coefficients were compared between different numbers of reference database communities based on Welch’s *t*-test in each experimental treatment.

### Community-level forecasting

The above population-level results based on empirical dynamics modeling were then used for forecasting community-level dynamics. For a focal time point (day) in a target experimental replicate, the 16S rRNA copy concentration estimates (predicted abundance) of respective ASVs were compiled, yielding predicted community structure (i.e., predicted relative abundance of ASVs). The predicted and observed (actual) ASV compositions (relative abundance) of respective target replicates were then plotted on a NMDS surface for each of the six experimental treatments. In addition, we evaluated dependence of community-level forecasting results on experimental conditions (source inocula and media), *α*-diversity (Shannon’s *H′*) of ASVs, and mean *β*-diversity against other replicates in a multivariate ANOVA model of predicated vs. observed community dissimilarity.

### Anticipating abrupt community shifts

We then examined whether indices derived from the energy landscape analysis and/or empirical dynamics modeling could be used to anticipate drastic changes in community structure.

In the framework of energy landscape analysis, we calculated two types of indices based on the estimated landscapes of microbiome dynamics (Fig. 3a). One is deviation of current community-state energy from the possible lowest energy within the target basins (hereafter, energy gap; Fig. 3a): this index represents how current community states are inflated from local optima (i.e., “bottom” of basins). The other is “stable-state entropy^24^”, which is calculated based on the random-walk-based simulation from current community states to bottoms of any energy landscape basins (i.e., alternative stable states). A starting community state is inferred to have high entropy if reached stable states are variable among random-walk iterations: the stable-state entropy is defined as the Shannon’s entropy of the final destinations of the random walk^24^. Therefore, communities approaching abrupt structural changes are expected to have high stable-state entropy because they are inferred to cross over “ridges” on energy landscapes. For an analysis of a target replicate community, energy landscapes were reconstructed based on the data of the remaining seven replicate communities.

In the framework of empirical dynamics modeling (nonlinear mechanics), we calculated “local Lyapunov stability^40^” (local dynamic stability) and “local structural stability^41^” based on Jacobian matrices representing movements around reconstructed attractors^27^. Specifically, based on convergent cross-mapping^22,32^ and multivariate extension of S-map^63^, local Lyapunov stability and structural stability were estimated, respectively, as the absolute value of the dominant eigenvalue and trace (sum of diagonal elements) of the Jacobian matrices representing the time-series processes^40^. For a target replicate community, the remaining seven replicate communities were used for inferring Jacobian matrices. Note that a high score of local Lyapunov/structural stability represents a potentially unstable community state. In particular, local Lyapunov scores reflect whether trajectories at any particular time are converging (local Lyapunov score < 1) or diverging (1 < local Lyapunov score)^40^.

For each of the above indices, linear regression of abruptness scores of community-compositional changes was performed for each replicate sample in each experimental treatment (seven-day-ahead forecasting). The time points (samples) excluded in the data-quality filtering process (see Bioinformatics sub-section) were excluded from this evaluation of signal indices.

We also examined the diagnostic performance of the signal indices based on the receiver operating characteristic (ROC) analysis. In seven-day-ahead forecasting, detection rates (sensitivity) and false detection rates (1 – specificity) of large community-compositional changes (abruptness > 0.5) were plotted on a two-dimensional surface for each experimental treatment, yielding area under the ROC curve (AUC) representing diagnostic performance^42^. The optimal threshold value of each signal index for anticipating abrupt community-compositional changes (abruptness > 0.5) was then calculated with the Youden index^42^ for each experimental treatment. In addition, for stable-state entropy and local Lyapunov stability, we calculated optimal threshold values after assembling all the data of Medium-A and Medium-B treatments.

## Data availability

The 16S rRNA sequencing data are available from the DNA Data Bank of Japan (DDBJ) with the accession number DRA013352, DRA013353, DRA013354, DRA013355, DRA013356, DRA013368 and DRA013379. The microbial community data are deposited at the figshare repository (DOI : 10.6084/m9.figshare.20653440).

## Code availability

All the scripts used to analyze the data are available at the figshare repository (DOI : 10.6084/m9.figshare.20653440).

## Acknowledgements

We thank Sayaka Suzuki and Keisuke Koba for support in the experiment and Tadashi Fukami for insightful comments on the manuscript. Computation time was provided by the SuperComputer System, Institute for Chemical Research, Kyoto University. This work was financially supported by JST PRESTO (JPMJPR16Q6), Human Frontier Science Program (RGP0029/2019), JSPS Grant-in-Aid for Scientific Research (20K20586), NEDO Moonshot Research and Development Program (JPNP18016), and JST FOREST (JPMJFR2048) to H.T., JSPS Grant-in-Aid for Scientific Research (20K06820 and 20H03010) to K.S., and JSPS Fellowship to H.F. and A.C‥

## Author Contributions

H.T. designed the work with H.F‥ H.F. performed experiments. H.F. analyzed the data with M.U., K.S., M.S.A., M.Y., and H.T‥ H.F. and H.T. wrote the paper with all the authors.

---

**Extended Data Fig. 1.**
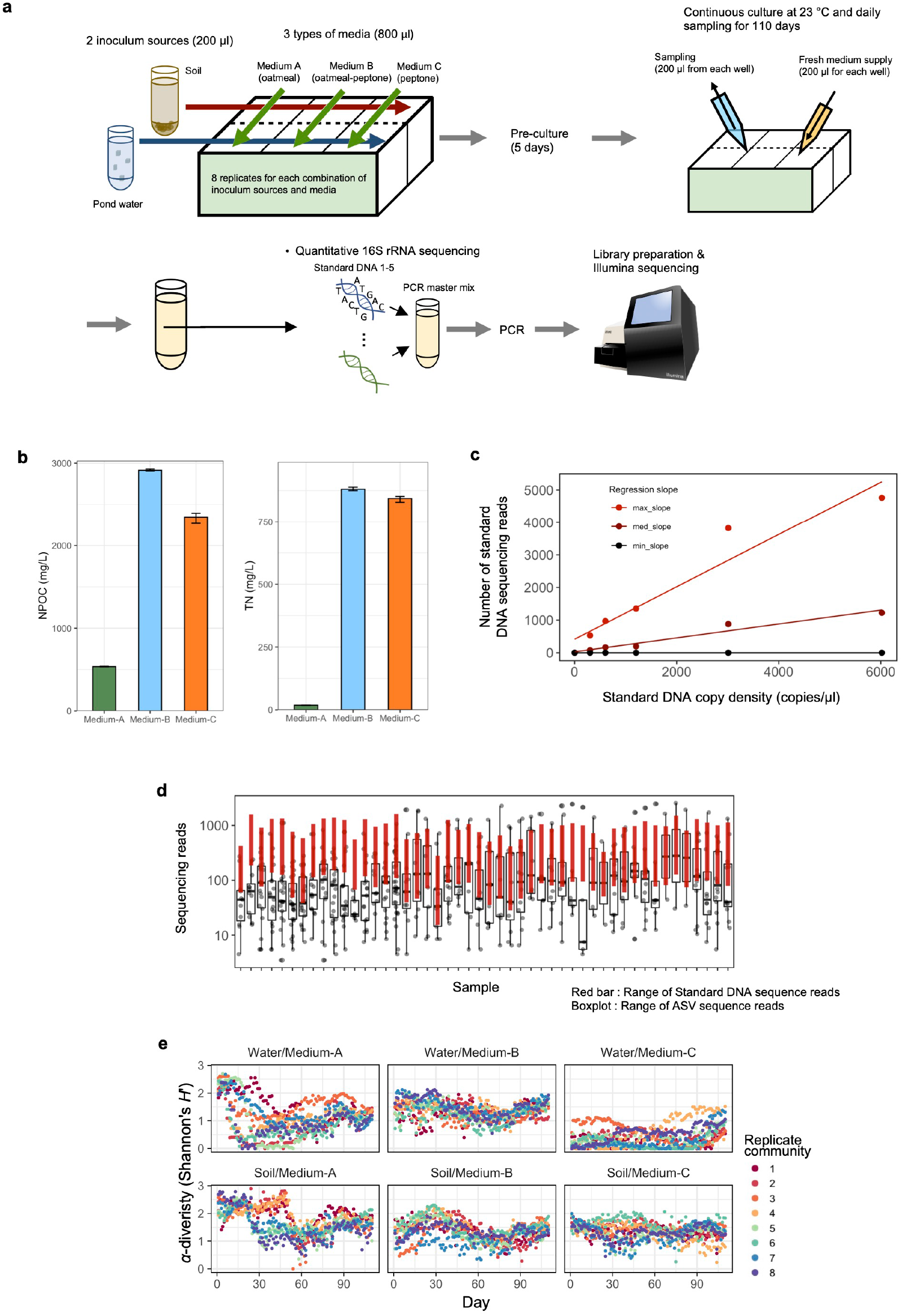
Experimental setting and microbiome data formats. **a**, Laboratoryculture system. Source microbiomes from forest soil and pond water were respectively introduced into three types of media [Medium-A, 0.5% (w/v) milled oatmeal; Medium-B, 0.5% (w/v) milled oatmeal + 0.5% (w/v) peptone; Medium-C, 0.5% (w/v) peptone] with eight replicates. A fraction of the culture fluid was sampled every 24 hours and equivalent volume of fresh medium was added to the continual culture system throughout the 110-day experiment. After DNA extraction, five “standard DNA” variants with different concentrations were introduced into the amplicon sequencing analysis of the 16S rRNA region, yielding DNA copy number estimates of each prokaryote ASV in each replicate sample. **b**, Concentrations of non-purgeable organic carbon (NPOC) and total nitrogen (TN) in each of the three types of fresh media. The bars represent ranges of triplicate measurements. **c**, Example of calibration of 16S rRNA copy concentration. In most microbiome studies, only proportions of respective microbe’s sequencing reads to total sequencing reads (relative abundance; Extended Data Fig. 3) have been analyzed, while calibrated abundance information (absolute abundance; Extended Data Fig. 2) allows us to reconstruct population dynamics (i.e., increase/decrease through time-series) of respective ASVs in microbiomes. Five standard DNA sequences varying in concentration were added to PCR master mix solutions to infer relationship between DNA copy concentration and the number of sequencing reads in each sample. **d**, Calibration of DNA copy concentration with the standard DNA gradients. The number of sequencing reads of each microbial ASV (boxplots and circles; black) was compared with that of standard DNA sequences (range of five standard DNA variants; red) in each sample. **e**, *α*-diversity (Shannon’s *H′*) of ASVs through the time-series.

**Extended Data Fig. 2.**
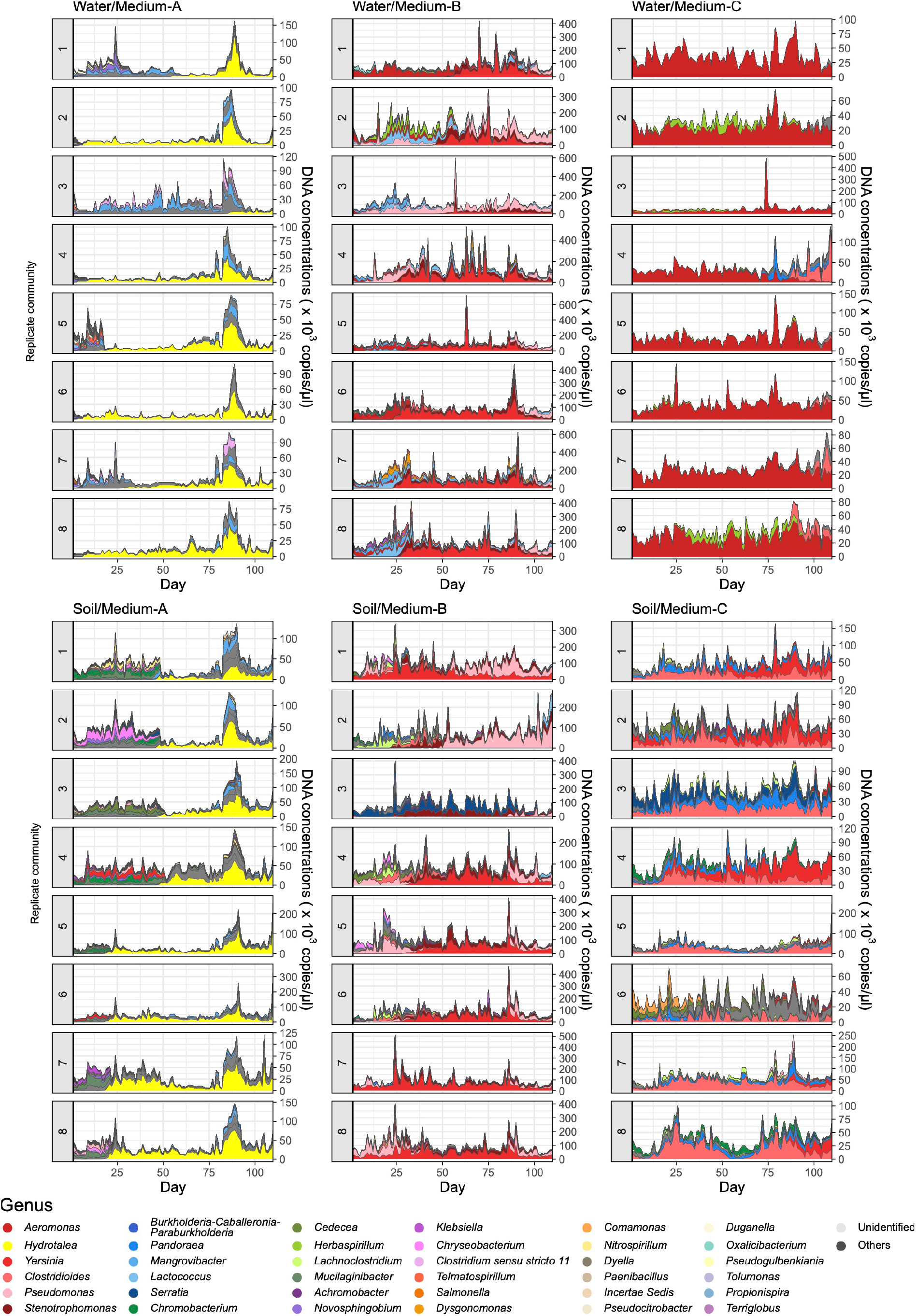
Dynamics of absolute abundance. For each replicate community of each experimental treatment, the changes of 16S rRNA gene copy concentrations (See Extended Data Fig. 1) are shown for each genus throughout the time-series. Note that each genus displayed in this figure can represent multiple microbial ASVs in the original dataset.

**Extended Data Fig. 3.**
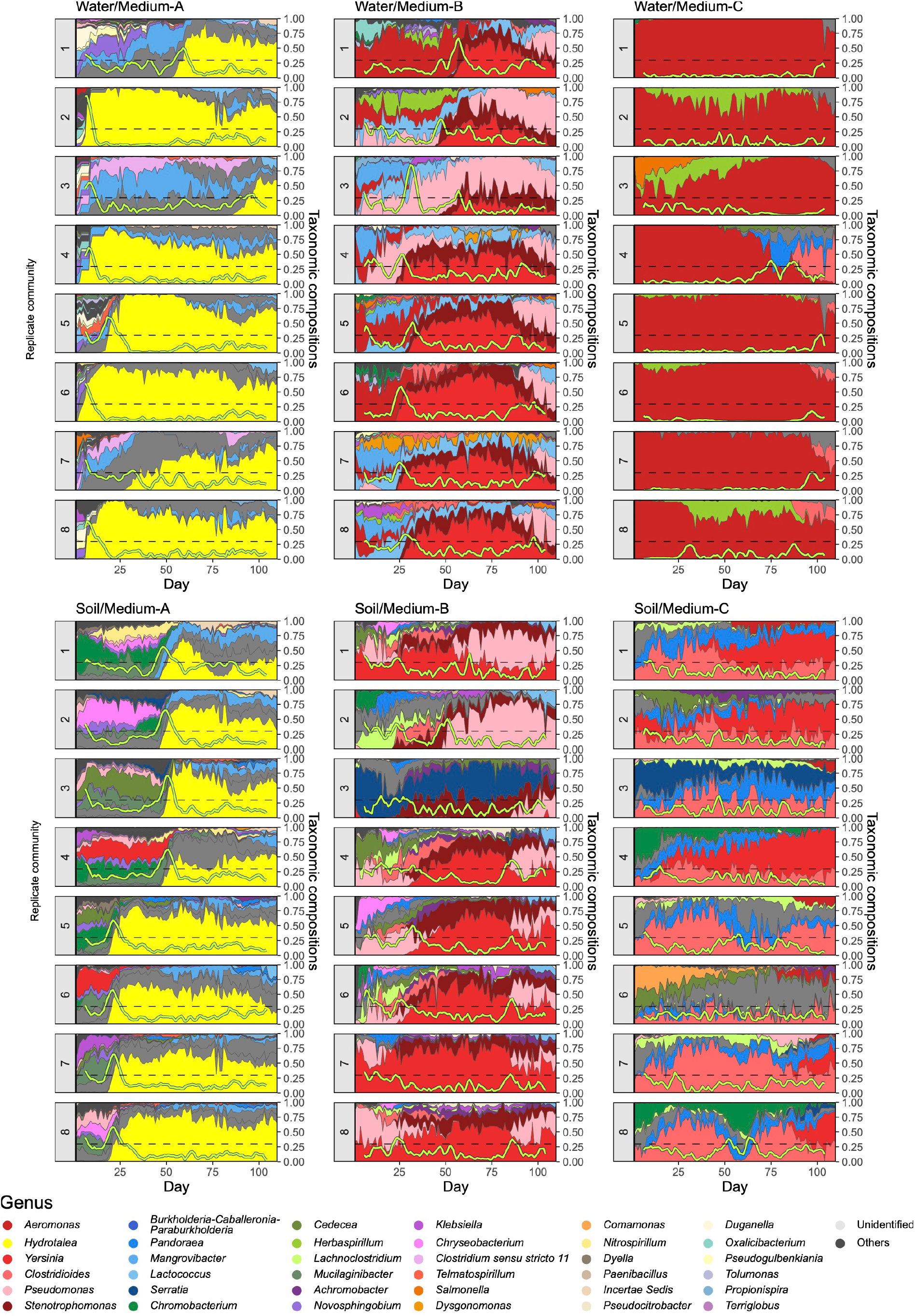
Dynamics of relative abundance. For each replicate community of each experimental treatment, the changes of the relative abundance of the 16S rRNA region are shown for each genus throughout the time-series. Note that each genus displayed in this figure can represent multiple microbial ASVs in the original dataset.

**Extended Data Fig. 4.**
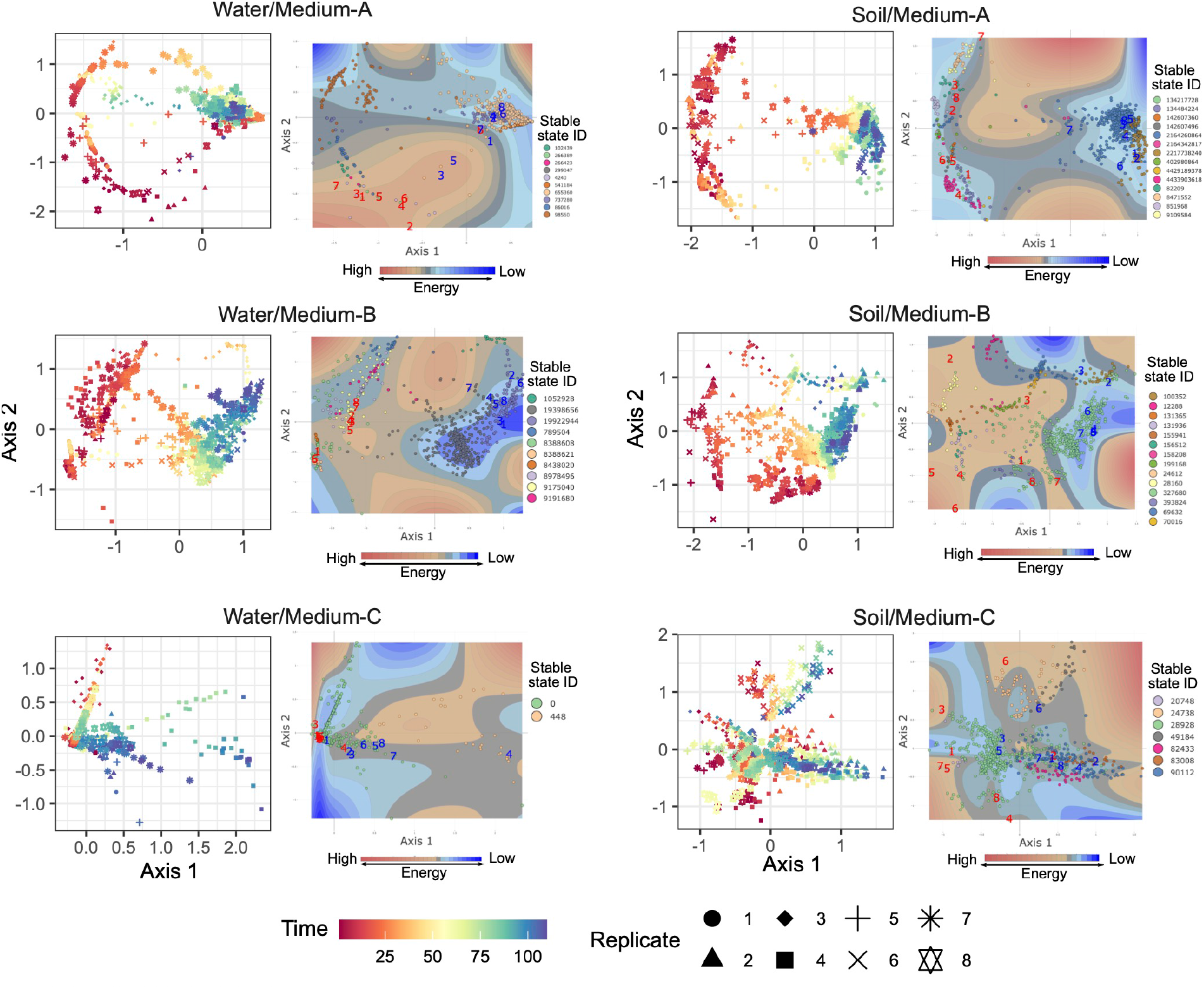
Distribution of stable states on the energy landscapes. The community structure of respective time points on NMDS axes (left) and reconstructed energy landscape on the NMDS surface (right) are shown for each experimental treatment. Community states (ASV compositions) located at lower-energy regions are inferred to be more stable on the energy landscapes. On the energy landscape of each experimental treatment, community states (data points) belonging to the basin of the same stable states are indicated with the same colors. The shapes of the landscapes were inferred based on a smoothing spline method with optimized penalty parameters. Within the energy landscape, community states of Day 1 and Day 110 are respectively shown in red and blue numbers representing replicate communities.

**Extended Data Fig. 5.**
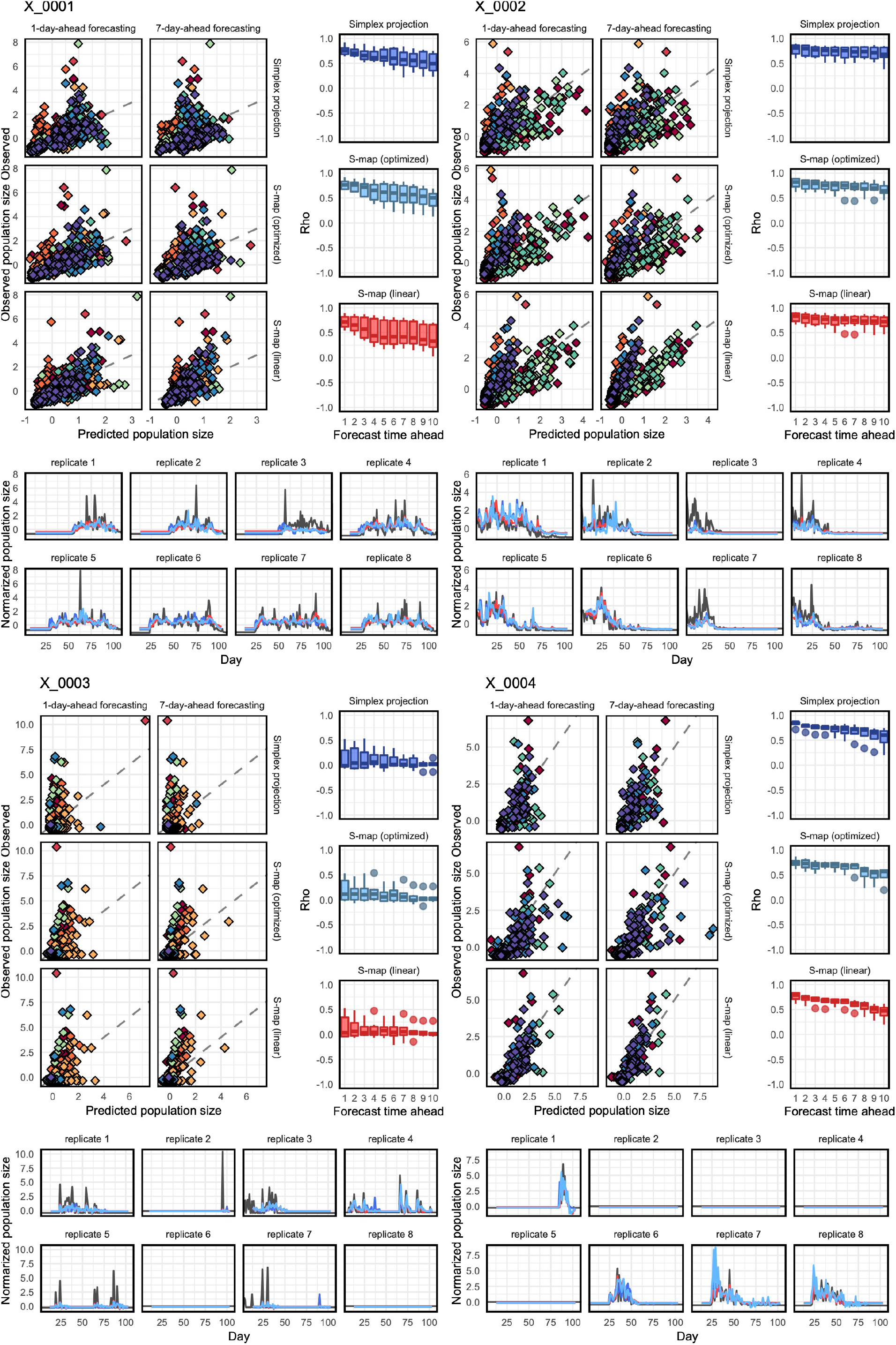
Examples of population-level forecasting results. For each microbial ASV in each experimental treatment, correlations between predicted and observed abundance through the time-series (one-day-ahead and seven-day-ahead forecasting; top left), decay of correlation between predicted and observed abundance (top right), and details of the time-series are shown. The prediction was based on simplex projection, S-map with optimized nonlinearity parameter (optimized *θ*), and S-map assuming linearity (*θ* = 0). For each target replicate community, the remaining seven replicate communities were used as references. Due to the large number of ASVs in the dataset, four ASVs in Water/Medium-B treatment are shown here as examples: the full results are available at the figshare repository (DOI : 10.6084/m9.figshare.20653440).

**Extended Data Fig. 6.**
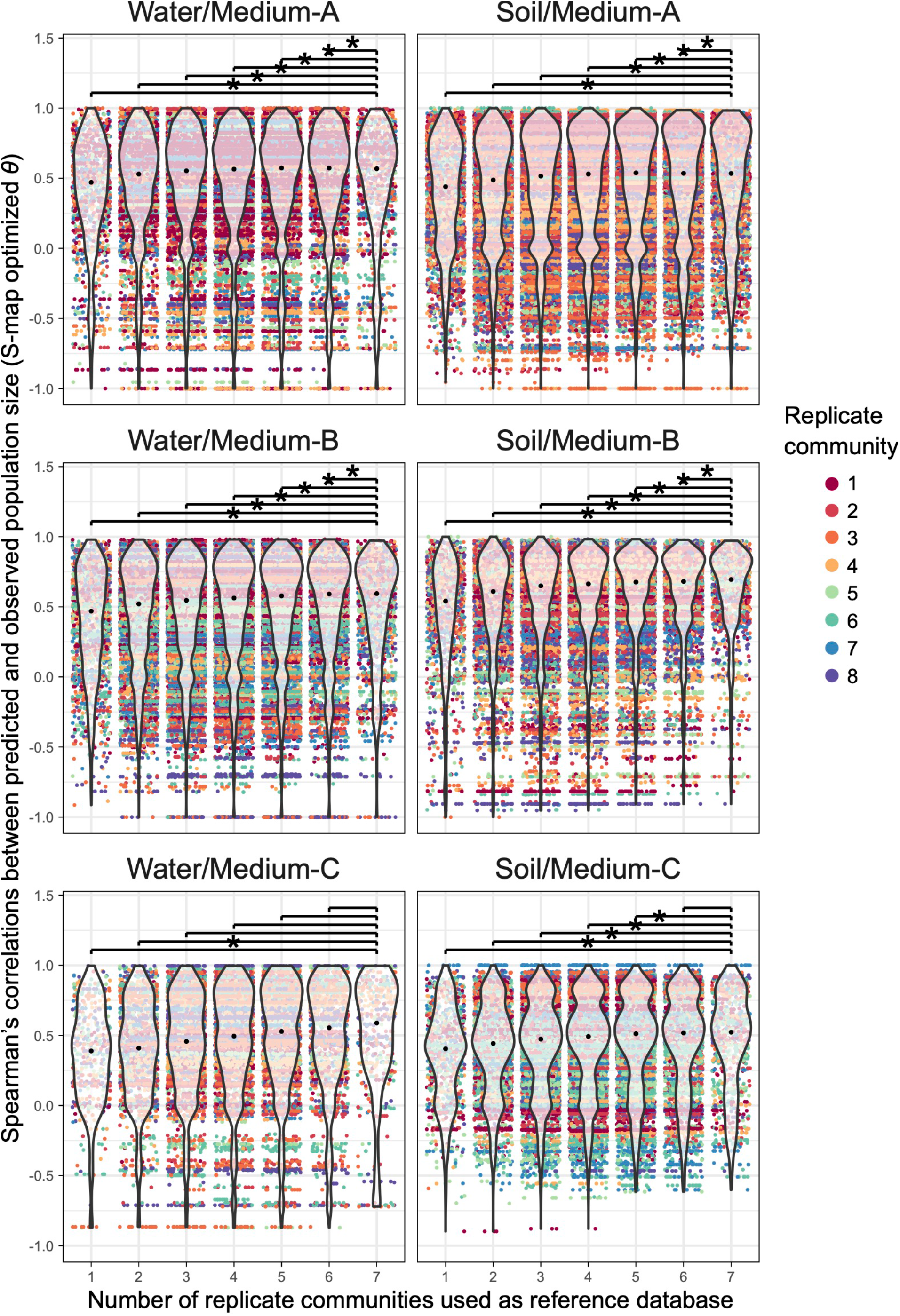
Dependence of population-level forecasting results on reference database size. The population size of each microbial ASV in a target replicate community was forecasted with S-map (optimized *θ*) based on reference databases (Fig. 2a). The forecasting was performed for each number of reference databases defined on the horizontal axis. Spearman’s correlations between predicted and observed population size (Fig. 2c) were calculated for each microbial ASV in each replicate community. An asterisk represents significant differences in forecasting skill (forecasting performance) between different numbers of reference databases in each experimental replicate: i.e., false discovery rate (FDR) based on Welch’s *t*-tests.

**Extended Data Fig. 7.**
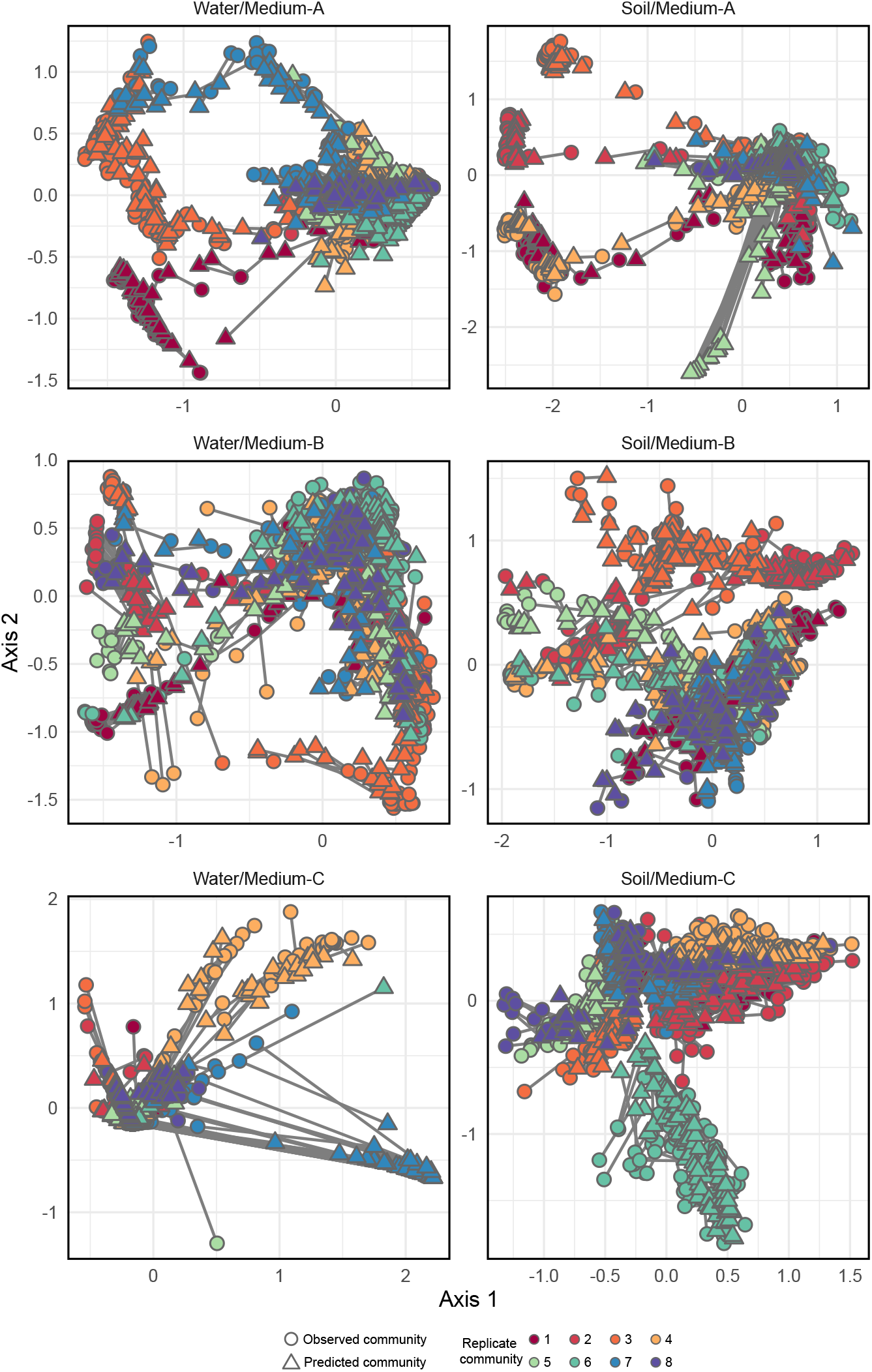
Comparison of predicted and observed community structure. By compiling the forecasting results of respective ASVs (Fig. 3; Extended Data Fig. 5), community compositions are predicted through the time-series. Predicted and observed community structure is linked for each day on the axes of NMDS (prediction based on S-map with optimized *θ*; one-day-ahead forecasting).

**Extended Data Fig. 8.**
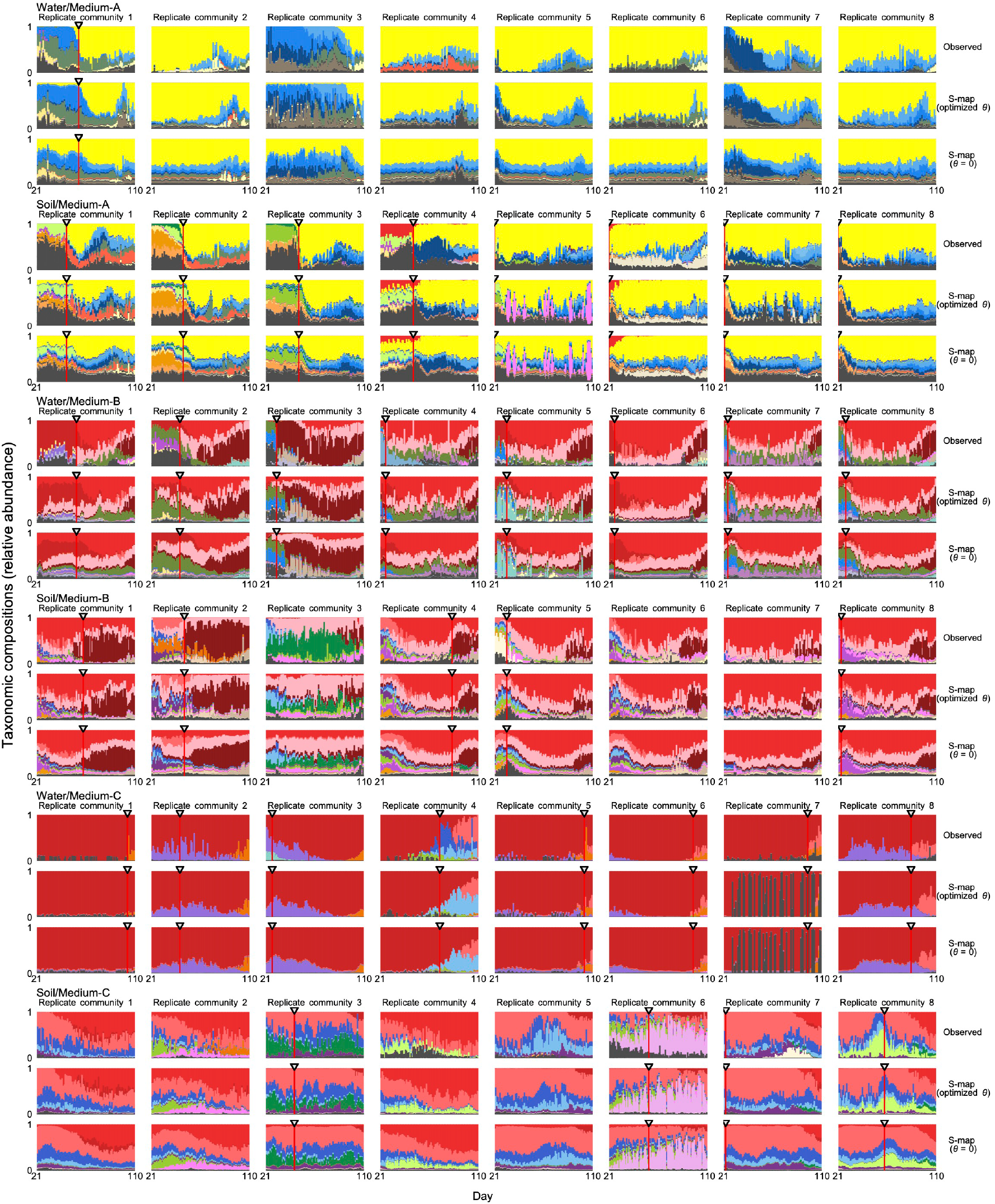
Comparison of nonlinear and linear forecasting approaches. Throughout the time-series, S-map nonlinear forecasting results are shown with observed community compositions and linear forecasting results (seven-day-ahead prediction). For the direct comparison of nonlinear and linear forecasting methods, S-map results with optimized nonlinearity parameter were compared with results of S-map assuming linear dynamics for all ASVs (*θ* = 0). Note that forecasting is inapplicable to the beginning of the time-series depending on embedding dimensions and forecasting time steps. A vertical line represents the timing of the greatest community compositional change in each replicate community.

**Extended Data Fig. 9.**
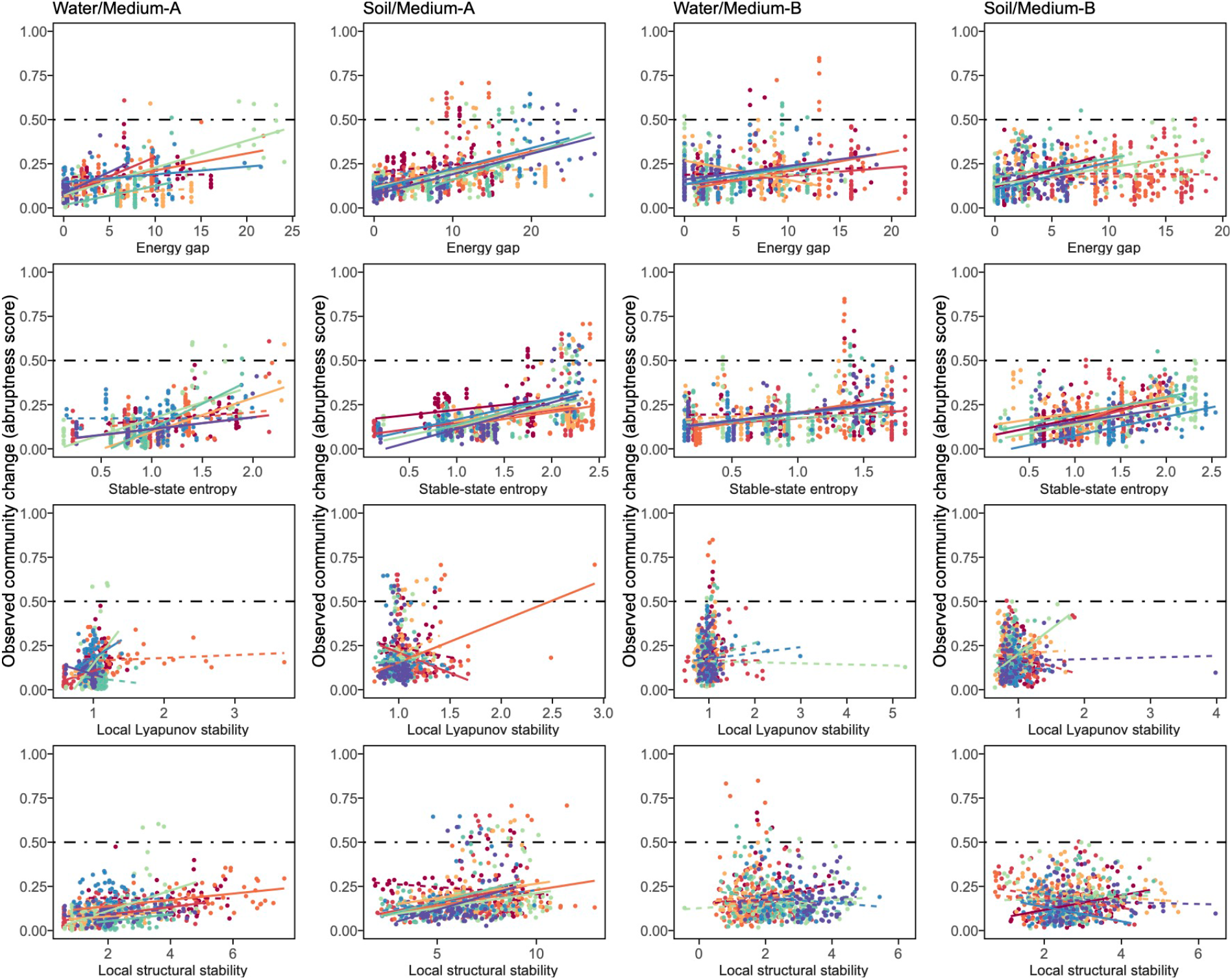
Candidates of signal indices for anticipating abrupt community changes. Relationships between signal index values and observed community-compositional changes are shown for seven-day-ahead forecasting. For each index of potential early-warning signals, Spearman’s correlation with the degree of community-compositional changes (abruptness scores) was examined for each time lag between signal indices and observed abruptness. The indices examined were the energy gap and stable-state entropy of the energy landscape analysis and the local Lyapunov stability and local structural stability of empirical dynamic modeling. Significant/non-significant regressions within respective replicates are shown with solid/dashed lines for each panel.

**Extended Data Fig. 10.**
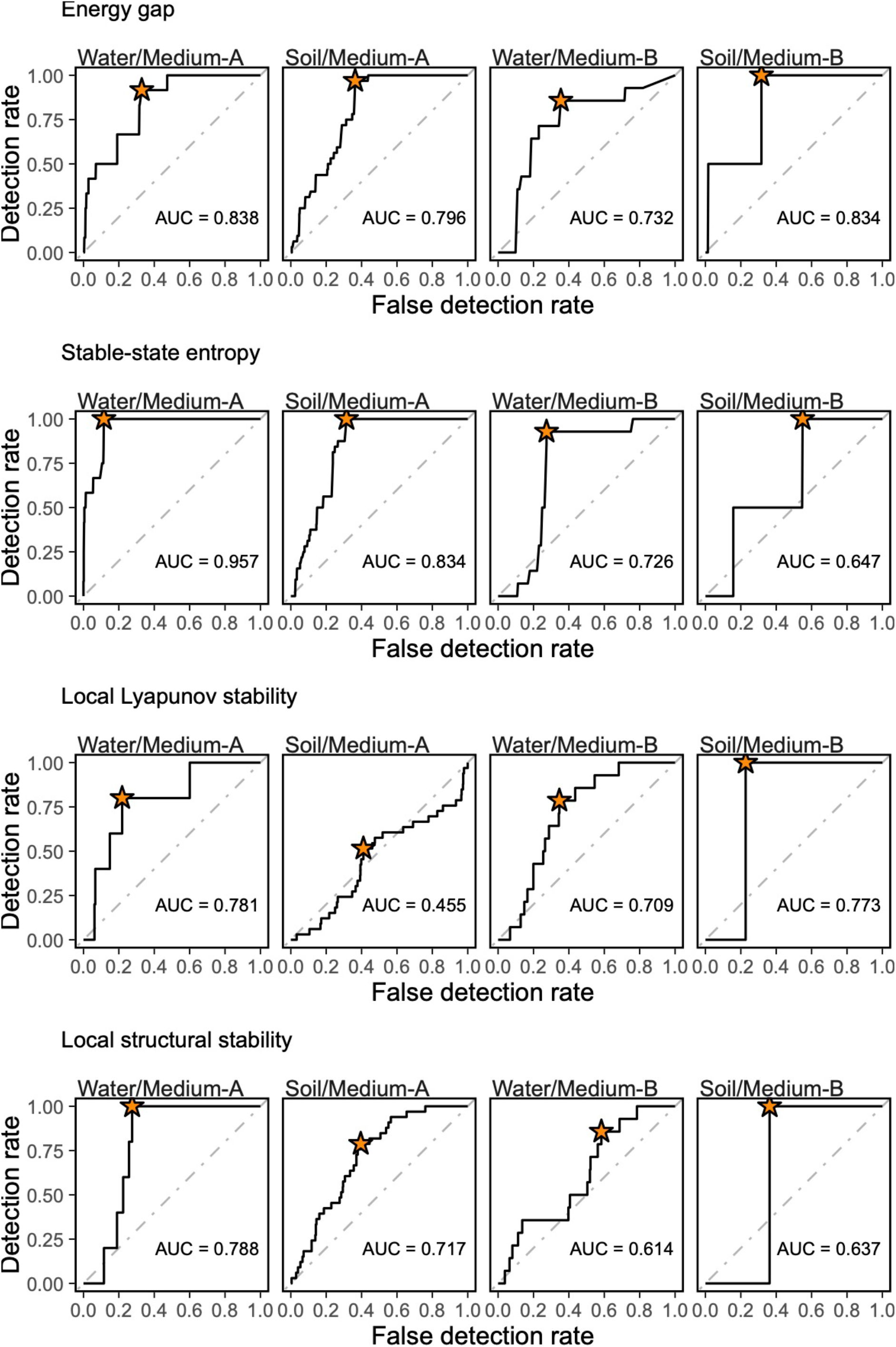
ROC analysis of diagnostic performance. On the two-dimensional surface of detection- and false-detection rates of abrupt community changes (abruptness > 0.5), area under the curve (AUC) and optimal detection rate (asterisk) were calculated (top panels) for local structural stability or energy gap. Optimal diagnostic threshold of local structural stability or energy gap for warning abrupt community changes was then obtained for each treatment based on the Youden index (bottom panels). Not that abrupt community changes were absent in Medium-C treatments and that the threshold for Soil/Medium-B treatment was unreliable due to the small number of time points with abruptness > 0.5 (Extended Data Figs. 3 and 9).

## References

1. Costello, E. K., Stagaman, K., Dethlefsen, L., Bohannan, B. J. M. & Relman, D. A. The application of ecological theory toward an understanding of the human microbiome. Science (1979) 336, 1255–1262 (2012).

2. Cho, I. & Blaser, M. J. The human microbiome: At the interface of health and disease. Nature Reviews Genetics 13, 260–270 (2012).

3. Huttenhower, C. et al. Structure, function and diversity of the healthy human microbiome. Nature 486, 207–214 (2012).

4. Wu, G. D. et al. Linking long-term dietary patterns with gut microbial enterotypes. Science (1979) 334, 105–108 (2011).

5. Toju, H. et al. Core microbiomes for sustainable agroecosystems. Nature Plants 4, 247–257 (2018).

6. Busby, P. E. et al. Research priorities for harnessing plant microbiomes in sustainable agriculture. PLoS Biology 15, e2001793 (2017).

7. Kazamia, E., Aldridge, D. C. & Smith, A. G. Synthetic ecology - A way forward for sustainable algal biofuel production? Journal of Biotechnology 162, 163–169 (2012).

8. Vatanen, T. et al. The human gut microbiome in early-onset type 1 diabetes from the TEDDY study. Nature 562, 589–594 (2018).

9. Zhao, L. et al. Gut bacteria selectively promoted by dietary fibers alleviate type 2 diabetes. Science (1979) 359, 1151–1156 (2018).

10. Kim, Y.-G. et al. Neonatal acquisition of Clostridia species protects against colonization by bacterial pathogens. Science (1979) 356, 315–319 (2017).

11. Chu, C. et al. The microbiota regulate neuronal function and fear extinction learning. Nature 574, 543–548 (2019).

12. Sato, Y. et al. Transcriptome analysis of activated sludge microbiomes reveals an unexpected role of minority nitrifiers in carbon metabolism. Communications Biology 2, 179 (2019).

13. Carding, S., Verbeke, K., Vipond, D. T., Corfe, B. M. & Owen, L. J. Dysbiosis of the gut microbiota in disease. Microbial Ecology in Health & Disease 26, 26191 (2015).

14. Ravel, J. et al. Daily temporal dynamics of vaginal microbiota before, during and after episodes of bacterial vaginosis. Microbiome 1, 29 (2013).

15. Hastings, A. et al. Transient phenomena in ecology. Science (1979) 361, (2018).

16. Fukami, T. Historical contingency in community assembly: integrating niches, species pools, and priority effects. Annual Review of Ecology, Evolution, and Systematics 46, 1–23 (2015).

17. Beisner, B. E., Haydon, D. T. & Cuddington, K. Alternative stable states in ecology. Frontiers in Ecology and the Environment 1, 376–382 (2003).

18. Hsieh, C. H., Glaser, S. M., Lucas, A. J. & Sugihara, G. Distinguishing random environmental fluctuations from ecological catastrophes for the North Pacific Ocean. Nature 435, 336–340 (2005).

19. Sugihara G. Nonlinear forecasting for the classification of natural time series. Philosophical Transactions of the Royal Society of London. Series A: Physical and Engineering Sciences 348, 477–495 (1994).

20. Sugihara, G. & May, R. M. Nonlinear forecasting as a way of distinguishing chaos from measurement error in time series. Nature 344, 734–741 (1990).

21. Benincá, E. et al. Chaos in a long-term experiment with a plankton community. Nature 451, 822–825 (2008).

22. Sugihara, G. et al. Detecting causality in complex ecosystems. Science 338, 496–500 (2012).

23. Strogatz, S. H. Nonlinear Dynamics and Chaos: With Applications to Physics, Biology, Chemistry, and Engineering. (CRC Press, 2015).

24. Suzuki, K., Nakaoka, S., Fukuda, S. & Masuya, H. Energy landscape analysis elucidates the multistability of ecological communities across environmental gradients. Ecological Monographs 91, 1–21 (2021).

25. Watanabe, T., Masuda, N., Megumi, F., Kanai, R. & Rees, G. Energy landscape and dynamics of brain activity during human bistable perception. Nature Communications 5, 4765 (2014).

26. Becker, O. M. & Karplus, M. The topology of multidimensional potential energy surfaces: Theory and application to peptide structure and kinetics. Journal of Chemical Physics 106, (1997).

27. Deyle, E. R., May, R. M., Munch, S. B. & Sugihara, G. Tracking and forecasting ecosystem interactions in real time. Proceedings of the Royal Society B: Biological Sciences 283, 20152258 (2016).

28. Chang, C. W., Ushio, M. & Hsieh, C. hao. Empirical dynamic modeling for beginners. Ecological Research 32, 785–796 (2017).

29. Munch, S. B., Brias, A., Sugihara, G. & Rogers, T. L. Frequently asked questions about nonlinear dynamics and empirical dynamic modelling. ICES Journal of Marine Science (2019) doi:10.1093/icesjms/fsz209.

30. Lahti, L., Salojärvi, J., Salonen, A., Scheffer, M. & de Vos, W. M. Tipping elements in the human intestinal ecosystem. Nature Communications 5, 1–10 (2014).

31. David, L. A. et al. Diet rapidly and reproducibly alters the human gut microbiome. Nature 505, 559–563 (2014).

32. Ushio, M. Interaction capacity as a potential driver of community diversity. Proceedings of the Royal Society B: Biological Sciences 289, 20212690 (2022).

33. Goldford, J. E. et al. Emergent simplicity in microbial community assembly. Science (1979) 361, 469–474 (2018).

34. Scheffer, M. & Carpenter, S. R. Catastrophic regime shifts in ecosystems: Linking theory to observation. Trends in Ecology and Evolution 18, 648–656 (2003).

35. Scheffer, M., Carpenter, S., Foley, J. a, Folke, C. & Walker, B. Catastrophic shifts in ecosystems. Nature 413, 591–596 (2001).

36. May, R. M. Thresholds and breakpoints in ecosystems with a multiplicity of stable states. Nature vol. 269 Preprint at https://doi.org/10.1038/269471a0 (1977).

37. Ovaskainen, O. et al. How are species interactions structured in species-rich communities? A new method for analysing time-series data. Proceedings of the Royal Society B: Biological Sciences 284, 20170768 (2017).

38. Hsieh, C. H., Glaser, S. M., Lucas, A. J. & Sugihara, G. Distinguishing random environmental fluctuations from ecological catastrophes for the North Pacific Ocean. Nature 435, 336–340 (2005).

39. Takens F. Detecting strange attractors in turbulence. in Dynamical Systems and Turbulence (eds. Rand DA & Young L-S) 366–381 (Springer, 1981).

40. Ushio, M. et al. Fluctuating interaction network and time-varying stability of a natural fish community. Nature 554, 360–363 (2018).

41. Cenci, S. & Saavedra, S. Non-parametric estimation of the structural stability of non-equilibrium community dynamics. Nature Ecology and Evolution 3, 912–918 (2019).

42. Akobeng, A. K. Understanding diagnostic tests 3: Receiver operating characteristic curves. Acta Paediatrica, International Journal of Paediatrics 96, 644–647 (2007).

43. Chang, C. et al. Reconstructing large interaction networks from empirical time series data. Ecology Letters 24, 2763–2774 (2021).

44. Amor, D. R., Ratzke, C. & Gore, J. Transient invaders can induce shifts between alternative stable states of microbial communities. Science Advances 6, 1–9 (2020).

45. Jones, C. G., Lawton, J. H. & Shachak, M. Organisms as ecosystem engineers. Oikos 69, 373–386 (1994).

46. Odling-Smee, F. J., Laland, K. N. & Feldman, M. W. Niche construction: The neglected process in evolution. Niche Construction: The Neglected Process in Evolution (MPB-37) vol. 9781400847266 (2013).

47. Mee, M. T., Collins, J. J., Church, G. M. & Wang, H. H. Syntrophic exchange in synthetic microbial communities. Proc Natl Acad Sci U S A 111, E2149–E2156 (2014).

48. Vrancken, G., Gregory, A. C., Huys, G. R. B., Faust, K. & Raes, J. Synthetic ecology of the human gut microbiota. Nature Reviews Microbiology 17, 754–763 (2019).

49. Ushio, M. et al. Quantitative monitoring of multispecies fish environmental DNA using high-throughput sequencing. Metabarcoding and Metagenomics 2, 1–15 (2018).

50. Caporaso, J. G. et al. Global patterns of 16S rRNA diversity at a depth of millions of sequences per sample. Proc Natl Acad Sci U S A 108, 4516–4522 (2011).

51. Apprill, A., Mcnally, S., Parsons, R. & Weber, L. Minor revision to V4 region SSU rRNA 806R gene primer greatly increases detection of SAR11 bacterioplankton. Aquatic Microbial Ecology 75, 129–137 (2015).

52. Klappenbach, J. A., Saxman, P. R., Cole, J. R. & Schmidt, T. M. Rrndb: The ribosomal RNA operon copy number database. Nucleic Acids Research 29, 181–184 (2001).

53. Lundberg, D. S., Yourstone, S., Mieczkowski, P., Jones, C. D. & Dangl, J. L. Practical innovations for high-throughput amplicon sequencing. Nature Methods 10, 999–1002 (2013).

54. Stevens, J. L., Jackson, R. L. & Olson, J. B. Slowing PCR ramp speed reduces chimera formation from environmental samples. Journal of Microbiological Methods 93, 203–205 (2013).

55. Hamady, M., Walker, J. J., Harris, J. K., Gold, N. J. & Knight, R. Error-correcting barcoded primers for pyrosequencing hundreds of samples in multiplex. Nature Methods 5, 235–237 (2008).

56. Tanabe, A. Claident v0.2.2018.05.29, a software distributed by author at http://www.fifthdimension.jp/. (2018).

57. Callahan, B. J. et al. DADA2: High-resolution sample inference from Illumina amplicon data. Nature Methods 13, 581–583 (2016).

58. Wang, Q., Garrity, G. M., Tiedje, J. M. & Cole, J. R. Naïve Bayesian classifier for rapid assignment of rRNA sequences into the new bacterial taxonomy. Applied and Environmental Microbiology 73, 5261–5267 (2007).

59. Quast, C. et al. The SILVA ribosomal RNA gene database project: Improved data processing and web-based tools. Nucleic Acids Research 41, D590–D596 (2013).

60. Oksanen, J. The vegan package available at https://cran.r-project.org/web/packages/vegan/index.html. (2007).

61. Wood, S. mgcv: Mixed GAM computation vehicle with automatic smoothness estimation available at https://cran.r-project.org/web/packages/mgcv/index.html. (2022).

62. Navarrete, R. Embeddings and prediction of dynamical time series. (The University of Michigan, 2018).

63. Cenci, S., Sugihara, G. & Saavedra, S. Regularized S-map for inference and forecasting with noisy ecological time series. Methods in Ecology and Evolution 10, 650–660 (2019).

